# Measure of major contents in animal and plant genomes, using Gnodes, finds under-assemblies of model plant, Daphnia, fire ant and others

**DOI:** 10.1101/2023.12.20.572422

**Authors:** Donald G. Gilbert

## Abstract

Significant discrepancies in genome sizes measured by cytometric methods versus DNA sequence estimates are frequent, including recent long-read DNA assemblies of plant and animal genomes. A new DNA sequence measure using a baseline of unique conserved genes, Gnodes, finds the larger cytometric measures are often accurate. DNA-informatic measures of size, as well as assembly methods, have errors in methodology that under-measure duplicated genome spans.

Major contents of several model and discrepant genomes are assessed here, including human, corn, chicken, insects, crustaceans, and the model plant. Transposons dominate larger genomes, structural repeats are often a major portion of smaller ones. Gene coding sequences are found in similar amounts across the taxonomic spread. The largest contributors to size discrepancies are higher-order repeats, but duplicated coding sequences are a significant missed content, and transposons in some examined species.

Informatics of measuring DNA and producing assemblies, including recent long-read telomere to telomere approaches, are subject to mistakes in operation and/or interpretation that are biased against repeats and duplications. Mistaken aspects include alignment methods that are inaccurate for high-copy duplicated spans; misclassification of true repetitive sequence as heterozygosity and artifact; software default settings that exclude high-copy DNA; and overly conservative data processing that reduces duplicated genomic spans. Re-assemblies with balanced methods recover the missing portions of problem genomes including model plant, water fleas and fire ant.

## Introduction

This paper describes measurements of the major contents of animal and plant genomes, comparing contents of chromosome assemblies with genomic DNA fragments used in their assembly. Four model organisms (human, corn, chicken, at-plant) are measured, Daphnia waterfleas and some other species that have difficult or discrepant genome assemblies. This paper focuses on large scale genomic content, and their discrepancies between assemblies and source DNA. Other content qualities at smaller scale are important and may contradict large scale effects.

Biochemical or cytometric methods of genome size estimation are well established, such as flow cytometry, and are subject to measurement errors of various kinds, but are not known to be biased (Dolezel & Greilhuber 2010, Pellicer & Leitch 2019). DNA sequence estimation methods, such as GenomeScope, are also subject to various measurement and parameter errors (Mgwatyu et al. 2020, Ranallo-Benavidez et al 2020).

There is often a difference in genome sizes measured from cytometry or DNA sequences, that DNA measures and assemblies are smaller than cytometric measures. Figure I1 illustrates this, comparing recent genome assemblies of animals and plants, versus their cytometric measured sizes, for about 100 species each of arthropods, birds, fishes, mammals and plants. A majority of assemblies are well below cytometric sizes, from 15% to over 50% smaller. There are many factors contributing to these discrepancies. This trend of smaller assemblies can be examined through measure of a third evidence, the major contents of the raw DNA sequence used to build these assemblies.

**Figure I1.**
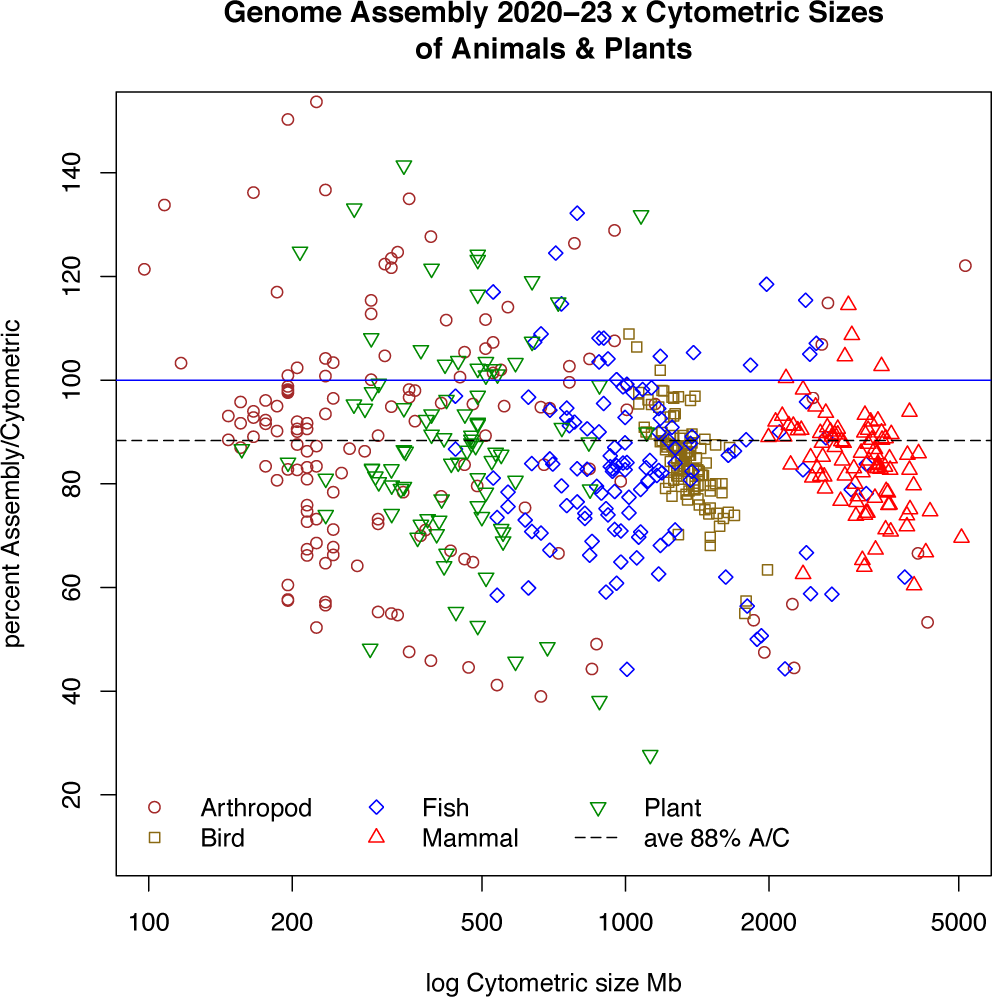
Genome sizes for 500 animal and plant species, as Assembly / Cytometric size percentage (Y) in relation to cytometric sizes (X-axis in log megabases), for approx. 100 species each of arthropods, birds, fishes, mammals and plants. NCBI Genomes (NCBI 2023) provides assembly sizes, median value for multiple assemblies, from year 2020 to 2023, many of long-read DNA methods. Animal Genome Size Database (Gregory 2023) and Plant DNA C-values Database (Leitch et al 2019) provide the most recent cytometric sizes of the same species [Suppl. table eukaryote_genoacsizes1a.txt].

Measurements are based on variants of the basic formula G= L*N/C (Lander & Waterman, 1988) to calculate genome size in bases (G) from observed DNA fragment sizes (L) and number of DNA reads (N), divided by a constant C coverage depth of those fragments. The C coverage value is found using unique conserved gene coding sequences as a reliable “inch” for a genome measurement ruler (Gnodes#1). For comparison, measurements are made using the method of K-mer genome estimations from peak of a poisson distribution of fixed-size oligomer sub-fragments of DNA fragments (Li and Waterman 2003).

The basic measurement method is to align, or map, DNA fragments with genome assemblies, where the assemblies have sequence contents annotated with coding genes, transposons, repetitive sequences and other items. Four major contents are measured, three are fairly reliably identifiable by their sequence patterns: 1. P, protein gene codes, 2. T, transposon codes, 3. R/S, repeated pattern sequences of major structural components (centro- and telomeres), and 4. O, the other, containing non-coding genes, introns and intergenic portions that lack those three patterns. There is some ambiguity in distinguishing organismal gene codes and transposon codes, which this paper does not try to resolve, thus there is overlap in these contents. Gnodes calculates amount of contents from coverage depths in DNA samples and assemblies, and copy numbers relative to unit unique gene sequences. The ratio of DNA sample to assembled genome, termed xCopy, indicates genome regions and contents that are under-assembled (xCopy > 1) or over-assembled (xCopy < 1) in simple cases.

An alternate measurement method also used here is to count all fixed-size (K-mer) oligomer sub-fragments of DNA samples and assemblies, and count oligomers of intersecting subsets for these contents. Sequence alignment methods are a common approach to quantifying genome contents, and allow for biological and artifact errors that exist. Perfect matching of K-mer fragments offers an alternate way to validate counts of very-high-copy duplicate sequences that are very difficult to accurately measure.

## Results of measuring major contents

Genome measurement results here are presented from large (human, 3000 megabase genome), through corn (2000 Mb) and chicken (1000 Mb) down to small arabidopsis model plant (150 Mb), the problematic Daphnia species (250 Mb by molecular measures, but assembled in several cases to half that), and other arthropods and plants with DNA/assembly discrepancies. This highlights the variability of major contents of genomes. Table 1 summarizes results for 12 species, models or where large discrepancies of cytometry versus assembly exist.

**Table 1.** Measures of genome size where large discrepancy exists.

| Species | Cyto size | DNA size | Assembly size Ref, Range | Content difference | Agreement |
| --- | --- | --- | --- | --- | --- |
| *Homo sapiens* | 3423 | 3130 | 3117, 3099-3117 | no | CYM > **DNA = ASM** |
| *Zea mays* | 2352 | 2293 | 2178, 2169-2179 | TE, CDS | **CYM = DNA** >= ASM |
| *Gal. gallus* | 1223 | 1210 | 1048, 1048-1058 | Other,CDS | **CYM = DNA** > ASM |
| *Lep. salmonis* | 1362 | 968 | 647, 643-647 | TE, HOR | CYM > DNA > ASM |
| *Sol. invicta* | 600 | 616 | 378, 374-478 | HOR | **CYM = DNA** > ASM |
| *Luc. sericata* | 675 | 535 | 565, 565-565 | no | CYM > **DNA = ASM** |
| *Luc. cuprina* | 616 | 520 | 409, 409-436 | HOR | CYM > DNA > ASM |
| *Dro. suzukii* | 328 | 308 | 268, 268-290 | LOR | **CYM = DNA** > ASM |
| *Dap. magna* | 240 | 237 | 161, 125-233 | HOR, CDS | **CYM = DNA** > ASM |
| *Dap. pulex* | 220 | 235 | 133, 133-227 | CDS, HOR,TE | **CYM = DNA** > ASM |
| *Dro. virilis* | 364 | 190 | 189, 182-189 | no | CYM > **DNA = ASM** |
| *Ara. thaliana* | 157 | 154 | 134, 119-146 | HOR,CDS | **CYM = DNA** > ASM |

Cytometric size: Flow cytometry or other cytometric genome size measure, median of values (CYM). DNA Size: DNA genome size measured with Gnodes (DNA). Assembly Size: Recent genome assembly sizes, reference (NCBI RefSeq) and range (ASM) Content difference: Main content differences for DNA > Assembly Agreement: How do Cytometric, DNA and Assembly sizes agree? Content types: CDS=coding sequences, TE=Transposons, HOR= higher-order-repeats or Satellite DNA, LOR= short simple repeats, Oth= Other or unannotated content;

A human genome has 47% Transposon codes, 40% Other, 11% R/Structure, and 2% Protein codes. A corn genome has 74% Transposon codes, 16% Other, 8% R/Structure, and 6% Protein codes. A chicken genome has 80% Other, 10% Transposon codes, 6% R/Structure, and 4% Protein codes. The 1/10th sized model plant genome has 30% Protein codes, 25% R/Structure repeats, 15% Transposons, and 30% Other. Figures M1, 2 and 3 plot these major contents for recent assemblies and DNA of nine in this taxonomic span, with cytometric sizes indicated. DNA measures up to cytometric sizes for 7 of 9, while assemblies are up to DNA levels for only 2 of the 9 species groups. There is probable biological variation among DNA samples; each comparison includes DNA used in an assembly plus other samples (see species details, and Suppl. table species23chrdna_gdupstr_sum1a.txt).

**Figure M1.**
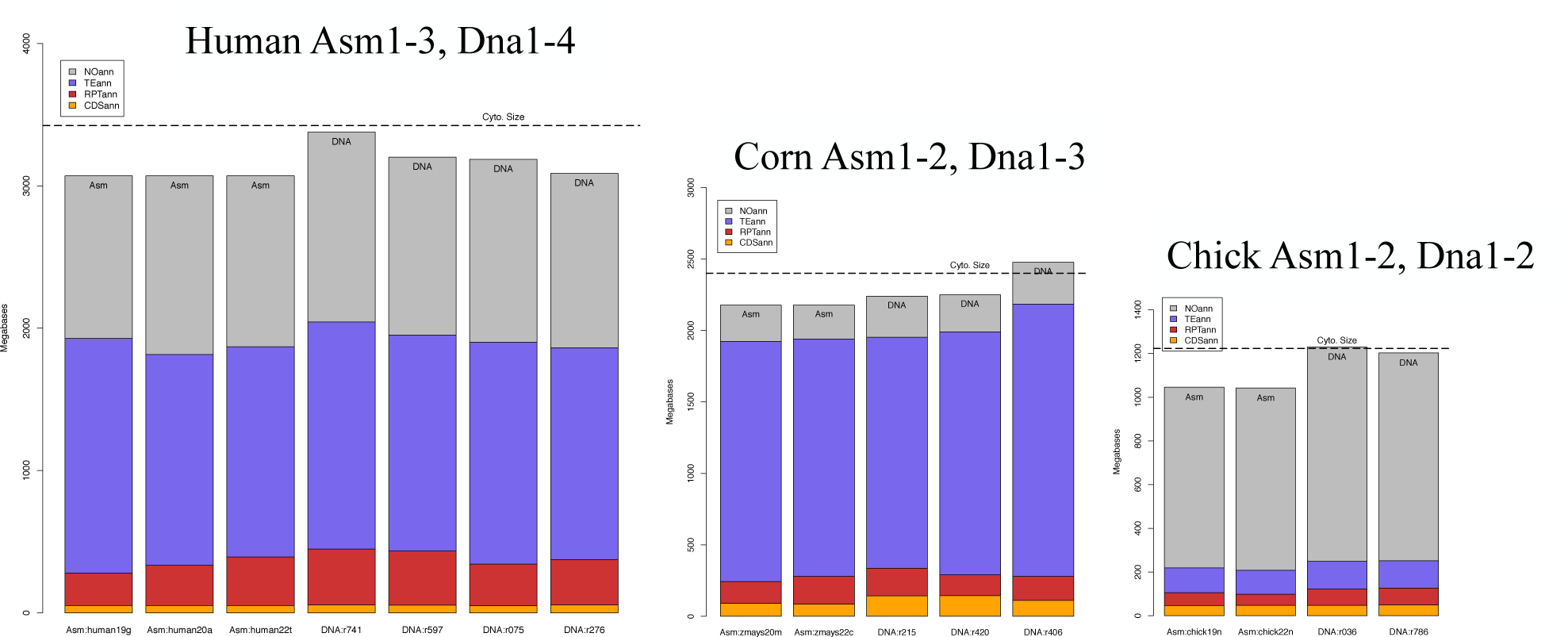
Major genome contents of human, corn, chicken genomes. Bars show megabase contents (Y-axis) of protein codes (orange, bottom), structural repeats (red-brown), transposons (blue) and other (gray), for several assemblies and genomic DNA samples (X-axis).

Insects and crustaceans with discrepant genome size measures (cytometric vs. sequence assembly) were examined, and cases with useful DNA samples are shown here. Tabulation of the discrepant sizes is made with Animal Genome Size Database (Gregory 2023) and NCBI Genomes assembly sizes (NCBI 2023) [Suppl. table asm_cytosize_table_arthropods.txt]. Published DNA samples corresponding to these are incomplete; among the discrepant with good samples are *Lucilia cuprina* and *L. sericata* (blowfly and bottle fly sibling species), *Solenopsis invicta* (red fire ant), *Lepeophtheirus salmonis* (salmon louse), and *Drosophila* flies.

**Figure M2.**
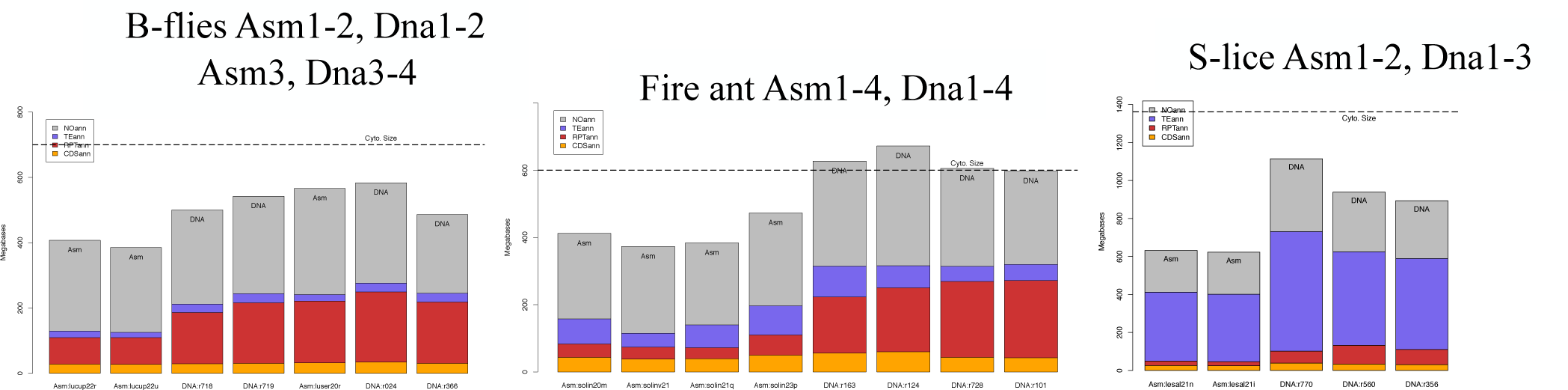
Major genome contents of insects blow flies, fire ant, and crustacean salmon louse genomes, content as per Figure M1

For Daphnia species, DNA contents measured in *D. pulex/pulicaria* are 45% Other, 26% Protein codes, 15-30% R/Structure, ad 12% Transposons, then D. magna has 35% Other, 27% Protein codes, 35-46% R/Structure, 12% Transposons. *D. magna* has more high-order, structural repeats by percentage of genome than any of these others, a factor complicating its assembly.

**Figure M3.**
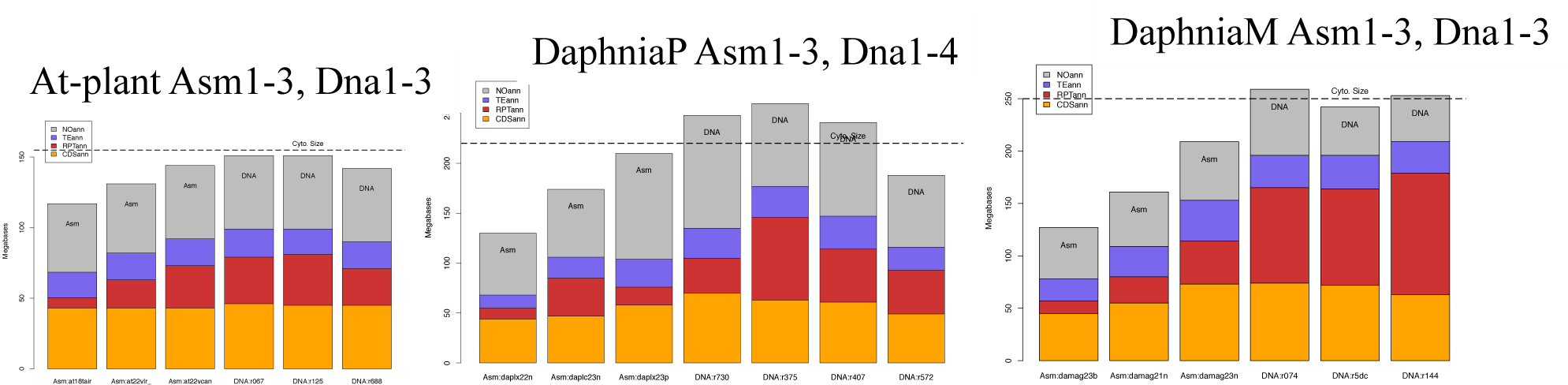
Major genome contents of small model plant and *Daphnia pulex/pulicaria* and *D. magna* genomes, content as per Figure M1.

Re-assemblies of genomic DNA produced for this paper are closer to the full contents measured in DNA, for *Arabidopsis*, *Daphnia pulex* and *magna, Drosophila suzukii,* and the fire ant *Sol. invicta.* These reassemblies all use long-read DNA sets (Pacbio, Nanopore) and Canu v2.2 assembler. Table 2 compares these re-assemblies with their reference assemblies of same DNA.

**Table 2.** Measures of genome size with large discrepancy, reference versus re-assembly of same long-read DNA.

| Species | DNA size | Re-asm | Ref. asm | Content difference | Notes |
| --- | --- | --- | --- | --- | --- |
| *Sol. invicta* | 616 | 478 | 378 | HOR | "Purge haplotigs" used on reference after assembly removed valid high order repeats |
| *Dro. suzukii* | 348 | 290 | 268 | LOR | Large spans of of short (5-10 bp), simple repeats, hard to assemble |
| *Dap. magna* | 237 | 233 | 161 | HOR, CDS | Largest portion of very high copy HOR, hard to assemble |
| *Dap. pulex* | 235 | 227 | 133 | CDS, HOR | Ref. is missing much of genome DNA duplications, due to unreported methods |
| *Ara. thaliana* | 154 | 146 | 134 | HOR | rDNA repeats at Chr2,4 telomeres are not fully captured in ref. |

DNA Size: DNA genome size measured with Gnodes (DNA).
Re-asm: Re-assembly of same long-read DNA as NCBI RefSeq assembly, using Canu
Ref. asm: Reference assembly size from NCBI RefSeq
Content difference: Main content differences for DNA > Assembly
Content types: CDS=coding sequences, TE=Transposons, HOR= higher-order-repeats or Satellite DNA,
LOR= short simple repeats, Oth= Other or unannotated content;
Notes: apparent causes for major content difference

The observed measures comparing DNA fragment and assembly contents indicate that these are often in agreement for both Transposon and Other categories (see chicken and corn for disagreement), across these and other plant & animal genomes examined. An inference is that these portions are divergent enough to assemble fully. In contrast, structural repeats, and to smaller extent coding sequences, are often not fully assembled where there is discrepancy in genome cytometric measures and assembly sizes. The genomic DNA contents contain more copies than are in assemblies, except for a few of the recent model genomes (Table 1). There are many animals and plants where cytometric and assembly sizes agree (Suppl. Table asm_cyto_size_equal_arthropods), not investigated here. Among the 60 *Drosophila* species with both measures, most agree. The discrepant cases find DNA agrees with assembly size but for *Dro. suzukii*. Of 5 *Drosophila* species examined for very high copy repeat content, all have simple repeats of 5-10 bases, in contrast to other species examined having longer (50-500 base) components of higher order repeats.

An overall summary of main content difference of genome assemblies and their DNA contents is shown in Figure T1. Few of these recent, mostly long-read assemblies reach the full contents found in their DNA, as indicated in Table 1. The largest contributor to this discrepancy is higher-order repeats, but also coding sequences are a significant missed content, and transposons for some. These long-read DNA sets can be more fully assembled, as Table 2 indicates, to capture this missed content. Human and corn plant genomes show the overall smallest percentage discrepancy, partly because their larger size leads the “mostly right” portion to dominate. Since many of these species were selected as having large discrepancies from cytometric measures, the result is expected.

**Figure T1.**
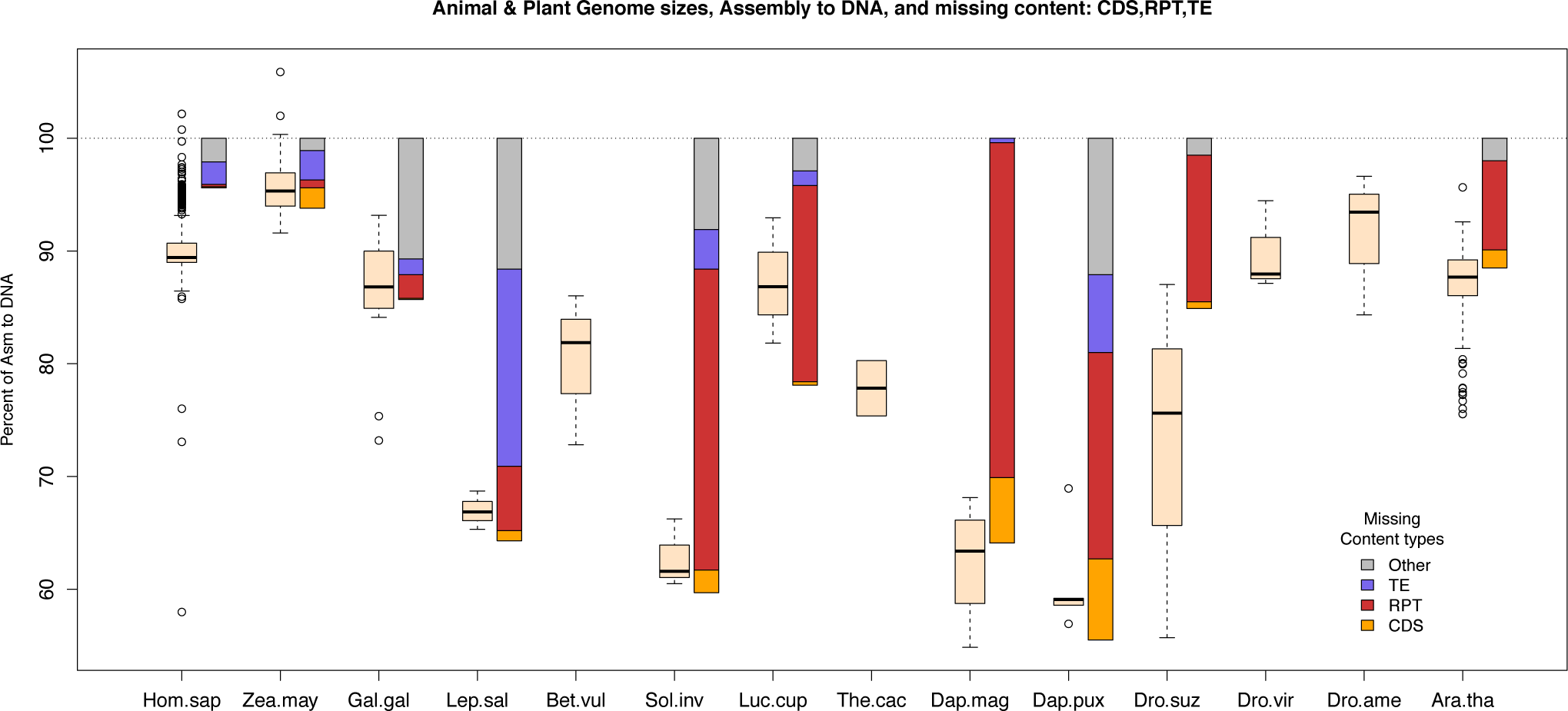
Animal and plant genome sizes, assembly relative to DNA contents, with missing major contents. Genomes are ordered big to small as in Table 1 on x-axis, with addition of *Beta_vulgaris* (sugar beet 743 mb), *Theobroma_cacao* (chocolate tree 431 mb), *Drosophila_americana* (190 Mb) that lack full repeat content analysis. Boxplot bars (bisque color) represent sizes from NCBI assemblies published at/after 2020, mostly long-read DNA, as percentage of the median measured DNA sizes on y-axis (60% to 100%). Missing contents of CDS, RPT, TE and Other are represented as Assembly/DNA percentage differences, stacked bars to right of genome boxplot, data as in figures M1,2,3. Data of suppl. table allspp_ncbiasmsize2020tab.txt

### Informatics problems with genomic repeats

Current informatics methods have difficulty assembling long, repetitive genomic spans with long-read DNA sequences. This should not be a surprise, but recent genome publications for many of the examined assemblies show little awareness of these problems. One value of Gnodes is to measure major contents of genomes in a way that is robust to high copy repeats and duplications, to provide evidence and reasons for discrepancies among size and content measures.

Several aspects of the informatics involved in producing genome assemblies, including recent telomere to telomere approaches, are subject to mistakes in operation and/or interpretation, which are often biased against repeats and duplications. These aspects include

a. reliance on alignment methods that are less accurate at high-copy duplicated spans;
b. mis-interpretation of true repetitive sequence as heterozygosity or artifact;
c. software default settings with high-end cut offs that exclude high-copy DNA;
d. unbalanced resolutions of ‘too-hot’ versus ‘too-cold’ problems in genomic data.

**Figure At23c4.**
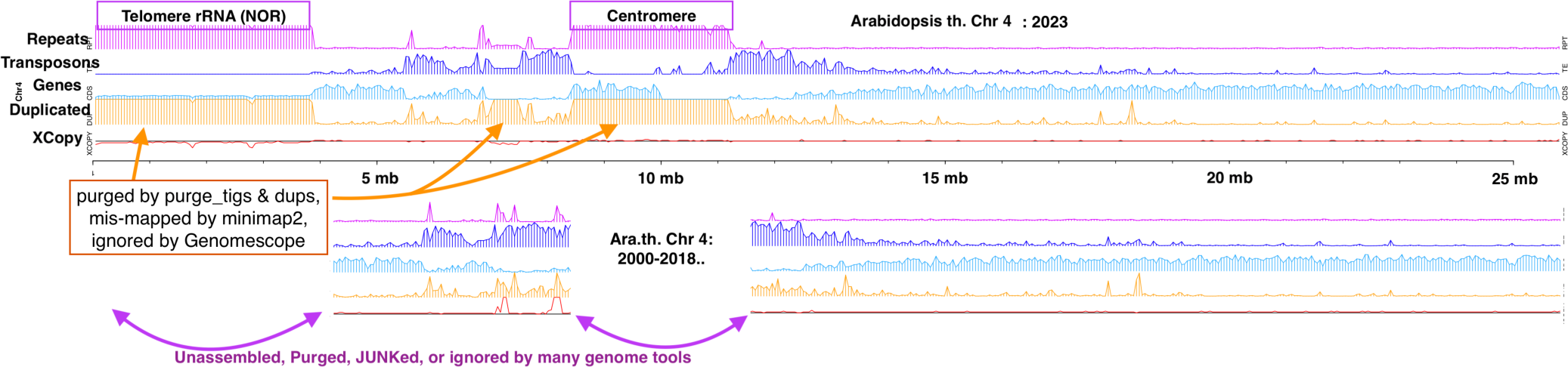
Example informatics problems with duplications shown in Gnodes main contents graph of Chromosome 4 of *Arabidopsis thaliana* (At, Col-0 strain), for new and reference assemblies, with annotations of genome software discrepancies. The top assembly, of 2023 this paper, includes a full nucleolus organizer region with highly identical, duplicated ribosomal RNA gene clusters, as well as a full centromere of high identity duplicated contents. The lower graph is current At reference assembly (TAIR10) that has not changed its major contents since its Sanger-sequenced publication (A.G.I., 2000). The discrepancy is the high copy, high identity duplicated spans. Many popular genome informatics tools are purging or ignoring those regions. See Gnodes#2 doc and below for details.

The model plant now has a complete genome assembly including its long repetitive telo- and centromeres, with a size that is 20% above what many genome informatics publications consider the “complete” model plant genome. This despite the fact that since the first At genome assembly (Arabidopsis Genome Initiative, 2000) it has been known to be incomplete; a portion of the chromosomes were found to be too repetitive to assemble, and that portion has only now, 23 years later, been added (Fultz et al. 2023, Naish et al 2021). At has a classic simple genome structure: mostly unique, euchromatic, gene-bearing arms of its five chromosomes, with repetitive centromeres midway, flanked by transposon-rich spans, and long repetitive telomeres on two chromosomes, containing nucleolus organizing regions. Figure At23c4 illustrates chromosome 4 of the model plant, the full assembly of 2023 and the partial assembly of 2000, updated through 2018 and still listed as the reference genome. Many genome software tools fail to process the long, duplicated spans of this example.

Measurements with Gnodes for both short and long read DNA, with several informatics tools, are reported in (Gnodes#2 doc). The short of it is that long and short read DNA will produce the same genome size and content measurements, with several caveats about biological, molecular and informatics methods used. This author found no fully reliable set of current (2022) methods for genome DNA measurements; any single method may produce very divergent results, versus a median of other methods. However a subset of DNA measurements is more likely to produce reliable results, ones that agree with either cytology measures or assembled DNA measures (Table 1).

#### A. Measuring/mapping high-copy duplications is difficult and inaccurate

Long-read DNA mapping introduces a problem of interpretation where parts of the long read do not map to an assembly, and other parts do. This is common in repetitive spans, where all bases of a long-read should be interpreted as belonging to a genome. Thus “unplaced” read bases are part of Gnodes analyses, which are placed for practical measures at the point of last mapping. With short-reads, unplaced bases are a minimal amount (1%). For long-reads these are significantly higher: 5% to 15%, and can be over 20% for some long-read data (fire ant HOR spans for example). These unplaced DNA spans, when extracted and re-aligned to their assembly, will mostly re-align at duplicated regions (90% mapped using blastn, but at lower rates using minimap2, for model plant, daphnia, fire ant and corn; Table 3A).

**Table 3A.**
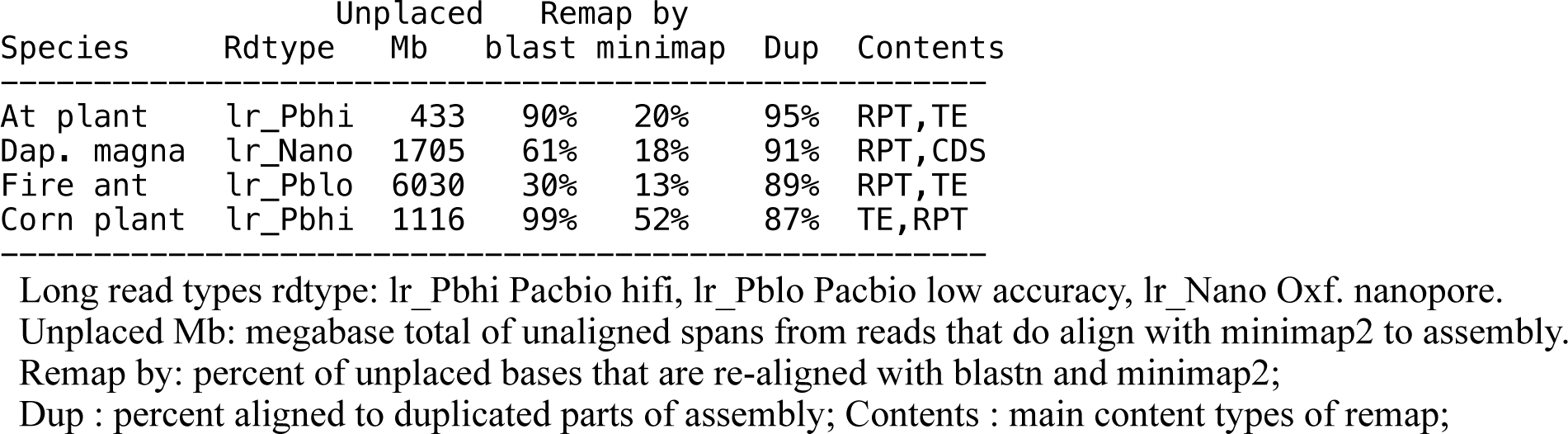
Re-map rates and contents of long-read unplaced spans; from Gnodes#2 table S7b_longread_unplacedbases.

The author has examined several long-read DNA mapping programs: minimap2 is a widely used now, and is used in Gnodes. Others that build on minimap2 include (1) Winnowmap2 (Jain et al. 2022), which adds k-mer analysis of repetitive DNA in mapping. Comparisons done for this project found winnowmap produces small differences from minimap2 which were inconsistent in terms of accurate measurement of repetitive content. (2) blend (Firtina et al. 2022), also based on minimap2, which its authors recommend for pacbio-hifi DNA. Tests for this project found that blend had lower hifi alignment compared to minimap, with more unplaced and unmapped reads. (3) lra (Ren et al. 2021), a long read aligner, which did map long reads, but poorly for this author; (4) TandemTools (Mikheenko et al. 2020), which failed to function for this author.

In general, error-full long read sequences (pacbio-lofi and oxford nanopore) are less accurately aligned and measured for coverage depth compared with highly accurate short-read DNA of current Illumina sequencers. This is a result of hundreds of Gnodes analyses across the range of animal and plant genomes. Winnowmap2 was designed to address this problem of inaccurate repeat mapping, and it is possibly a more accurate approach, with mistakes in this author’s tests of it (Gnodes#2, table S7c_read_mappers_long_short). This general inaccuracy in duplicate mapping is the reason that k-mer perfect match analyses of DNA x assembly copy numbers is added here to Gnodes analyses. While this second measure has some difficulties, results are similar enough with DNA x assembly alignments that both appear to be measuring the same underlying genomic components.

#### B. True duplications look like heterozygosity or artifacts, to some tools

This is a basic problem with genome data, that different contents can look the same with a limited range of measurement methods. Natural variation among duplicate sequences of different loci has similar signals to machine error and same-locus heterozygosity. For example, Jaegle et al. (2023) find that most “heterozygous” classed SNPs in *A. thaliana* are artifacts of duplicated regions, rather than residual heterozygosity. Genome informatics tools that look for one kind of content will mistakenly classify other kinds, when those tools don’t measure enough aspects to distinguish content kinds. Many tools that attempt to measure heterozygosity rely on alignment of DNA, and have problems noted in (A). One approach to resolving this is to classify contents that have different attributes or signals: repeats versus uniques, coding versus transposon versus satellite-type repeats, then apply measures for same-locus heterozygosity or artifacts, with different parameters suited to each class of sequence.

**Table 3B.** Removal of true duplications by popular heterozygosity dedup software, purge_dups and purge_haplotigs, for human and At plant on chromosomes with low and high Satellite DNA content.

|  |  | Size<br>Mb | SatDNA<br>Content | pur_dup<br>Removes | pur_tig<br>Removes |
| --- | --- | --- | --- | --- | --- |
| At_plant | Chr1 | 33 | 11% | 2% | 3% |
|  | Chr2 | 28 | 30% | 13% | 10% |
| Human | ChrX | 154 | 6% | 3% | 17% |
|  | ChrY | 62 | 45% | 59% | 65% |
|  | Chr8 | 146 | 5% | 3% | 4% |
|  | Chr9 | 151 | 23% | 12% | 19% |

At\_plant : at23vc2nl6b\_nor\_chrs split into 50kb parts, DNA of At Col0 Pbhifi SRR16841688 Human : human T2T chrs split to 50kb parts, DNA of HG002 son Pbhifi SRR18244889+890, in this male DNA, ChrX,Y are 1c, Chr8,9 are 2c

purge_haplotigs (Roach et al 2018), and the similar purge_dups (Guan et al 2020), are examples of tools designed to classify and remove assembly parts as spurious half-parts (alternate haploid spans assembled from a diploid genome). However, both misclassify true repetitive spans due to basic flaws (Table 3B): (1) mapping duplicated DNA to measure coverage, to identify “too-cold” low coverage and “too-hot” high coverage spans, fails because high copy, high identity repeats are poorly mapped, some repeat spans have under-counts, others have over-counts; (2) locating peaks (1x, 2x coverage) without careful coverage histogram analysis can produce false diploid/haploid classes; (3) classification as “junk”, which is no longer a valid genome data type, for many high copy repeats.

#### C. Improper defaults: e.g., truncated k-mer histograms

Default options for several genome tools are improper for measuring or retaining high-copy duplications. One reason for this is lack of software testing on known high-copy repeat genomes. This should change with recent reference assemblies, such as human Y-chr and model plant with full centro- and telo-meric repeat spans. Tools for assembly of unknown genomes should be validated on such now-known high-copy assemblies.

Jellyfish and other K-mer counters that produce the histograms used by GenomeScope, FindGSE, etc. have a default setting that limits output to 1,000 or 10,000 most unique kmers. This truncates all the high-copy repeated kmers, and thus truncates GSE estimates to mostly unique DNA. When a genome has portions of very high copy repeat DNA, these are missed. For genome size estimation with flow cytometry and k-mer analyses, Mgwatyu et al. (2020) found “.. GenomeScope [version 1] estimates were strongly affected by parameter settings, specifically CovMax. When using the complete k-mer frequency histogram, the programs did not deviate significantly [from flow cytometry size]”. False genome size estimates from truncated k-mer histogram results are replicated here (Table 3C), for model plant, *Daphnia magna*, *Sol. invicta*, and *Zea mays*. Full k-mer estimates and Gnodes estimates are in agreement with cytometry measures of genome size, while truncated histograms all under-estimate genome sizes.

**Table 3C.**
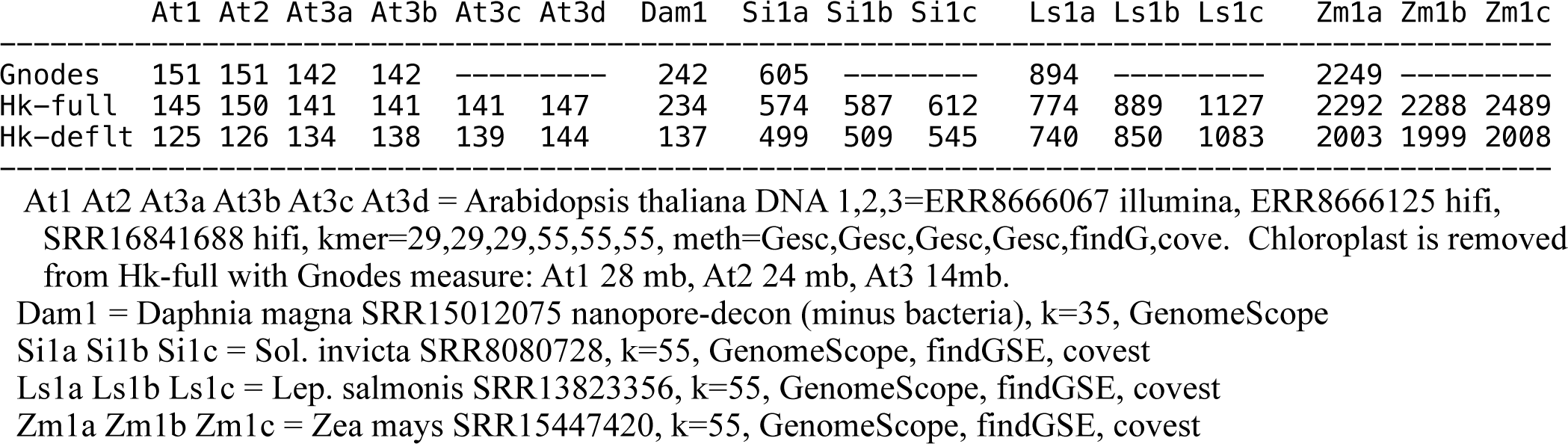
Genome size estimates from histograms of k-merized DNA samples, full (Hk-full) and default truncation at frequency 10,000 (Hk-deflt), for At plant, Daphnia magna, fire ant, salmon louse and corn plant. Sizes are estimated from histograms by GenomeScope, findGSE, covest-repeats. Gnodes estimate from DNA sample is also shown.

High-copy non-nuclear DNA, eg. plant chloroplasts, is included in full histogram measures, while truncated histograms truncate this along with high-copy nuclear DNA. This table adjusts GSE from full histograms using Gnodes-measured contaminant sizes (significant only for chloroplast; minor for mitochondria, bacteria). K-mer size choice has an effect on measuring repeats with truncated histograms, in that small K-mer values will lump more high-copy variants together, truncating more of these, but not or less effect for full histograms. Jellyfish and other K-mer calculators have options to set maximum counts in histogram; Jellyfish “histo --high 999999999” will produce a full histogram.

*TANSTAAFL in genomics*: In version 1 of GenomeScope (Vurture et al 2017), the authors explicitly add a truncation at coverage maxima of 1,000 as “those often represented organelle sequences or spike-in sequences occurring hundreds to thousands of times per cell in *A. thaliana* that artificially inflated the genome size”. This truncation does cut the DNA of 10,000 copies of chloroplast organelles found in some plant samples, but also cuts the high copy nuclear genome duplications of all samples. In version 2 of GenomeScope (Ranallo-Benavidez, Jaron & Schatz 2020), this problematic cut-off is removed, with a note in discussion that has been missed by many customers: “In addition, in order to accurately estimate genome size for highly repetitive genomes, **it is important to create a k-mer histogram that is not truncated**. By default, KMC and Jellyfish truncate the histogram at 10,000. We suggest running these tools without a maximum counter.” [bold added here].

## Discussion & Conclusions

### D. “This porridge is too cold”, complains Goldilocks

Genome informatics methods that reduce one type of error (spurious, extra assembly parts or sequence artifacts), often introduce the other type of error (removal of true duplications), due to lack of balanced evidence measurements. Overall, current genome informatics methods err on side of removing more true duplications than retaining them, sometimes with large imbalances to under-assembly (cf. *Daphnia magna* and *Dap. pulex*, fire ant examples). Genome assembly workflows or pipelines often lack an important engineering step: negative feedback tests to measure effects of each component on the overall evidence for assembly. E.g. steps that measure heterozygosity and remove putative heterozygous regions should also measure effects on high copy duplicates. The recently completed assembly of human Y chromosome (Rhie at al. 2023), rich in HOR or satellite DNA, has worked through these too-hot/too-cold trade-offs for a balanced approach.

Gnodes #1 documented under-estimates from GenomeScope and other k-mer measures, based on truncated histograms, as this author, like many peers, missed the very important detail about a bad default setting of genome tools. An important lesson for all genome projects and informatics tool developers, is to not rely on one method, but use two or more, with different underlying assumptions and algorithms, to resolve important aspects of genomes. Thus Gnodes and k-mer based measures of genome size and contents are complimentary, using different approaches to measure the same biological contents. With both approaches, a frequent disagreement of genome size estimates by cytometry and k-merized DNA estimates has been resolved, often but not always in favor of the larger cytometry measures (Tables 1, 2 and Figure T1 above).

### Satellite DNA in genome assembly problems

Satellite DNA, or higher-order repeats (HOR), are long DNA spans composed of shorter, common repeats, with variation. SatDNA covers large spans of centromeres, telomeres and interspersed regions of animal, plant genomes. In model plant, the recently assembled centromeres and telomeric NORs are higher order repeats of essential genes and structural DNA in multi-megabase spans for roughly 25% (37 mb) of an average At plant genome. In *Daphnia magna,* SatDNA appears to be as much as 40% of this genome, but most assemblies of this species have left out most of this content (e.g. only 4% in damag23bham, 10% in damag21ni7ncbi). In other *Daphnia* species, and across populations of species, SatDNA content is quite variable. Sibling species *D. pulex* and *pulicaria* vary from 3% to over 12% (based on under-assemblies, not DNA). In the fire ant, 20-30% of genome appears to be higher order repeats, mostly missing from current assemblies [see Fig. T1]. A recent report on *Tribolium* beetle measures SatDNA at 25% of 200 Mb genome (Marin et al 2023).

At 20% to 40%, SatDNA is a rather large portion of model plant, fire ant and *Daphnia* genomes, relative to other examined species. Such tandem repeats are very difficult to measure accurately, let alone assemble, even with longest read DNA methods. These numerous, large repeat spans confuse the assembly of nearby gene-bearing regions as well. Humans have approx. 7% SatDNA, corn and chicken also have a relatively smaller portion as SatDNA. It may be that HOR spans are not greatly expandable in larger genomes, compared with the expanded, self-replicating transposon content.

### Conclusions

Genome sizes from assemblies of animals and plants are often smaller than cytometric measures. A third measure, from DNA sequence samples before assembly, divided by the unit depth of conserved unique genes, generally support the larger cytometric measures. The major contents identified by this third measure indicates that structural, higher-order repeats are often under-assembled or missing from recent chromosome assemblies. This is a variable problem, small in many organisms but large in some, and it is amenable to measurement. Several commonly used genome informatics methods and tools contribute to the missing content by under-counting or mis-interpreting tandem and other genomic duplications. These under-assembly problems can be aleviated by use of balanced approaches to genome measurement and assembly.

## Example Species Details

### Human

**Figure Hs1a,b.**
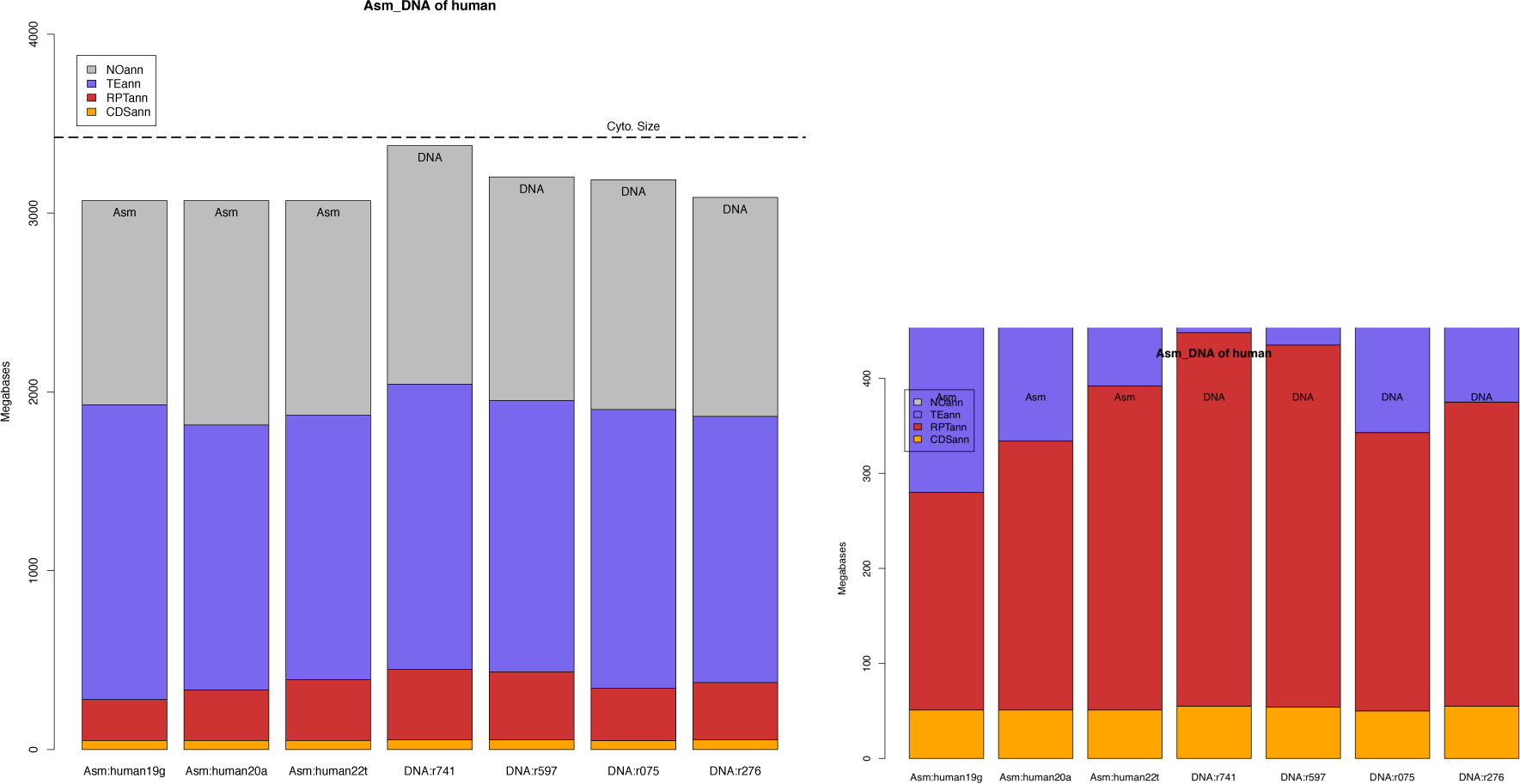
Major contents of human genome assemblies and DNA read sets measured with alignment of DNA on assemblies. Left panel (1a) has stacked bar graph of protein codes (orange, bottom), structural repeats (red-brown), transposons (blue) and other (gray) as portion of the 3100 megabase human genome. Three assemblies on left (2019 GRC, 2020 “Ash”, and 2022 T2T) are nearly same size, but with different contents. Four DNA read sets on right, of long and short-read form, are described in text. Right panel (1b) is a zoom down to 400-megabase bottom portion of contents to illustrate coding and structural repeat variation, comparable with smaller genomes.

**Figure Hs1c.**
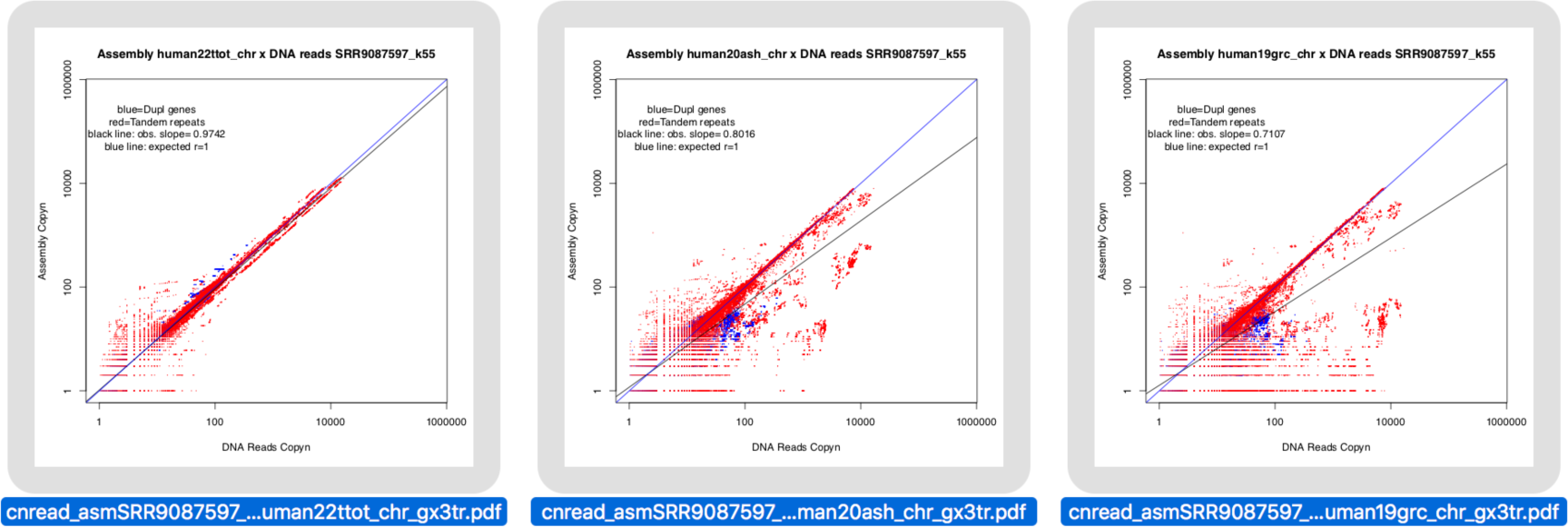
Perfect match counts of 55-mers of high-copy DNA by Assembly for human, for gene and structural content subsets. These plot the frequency of mers in bi-variate histograms, at the intersections of DNA and assembly contents. The X-axis is copy number in one DNA read set, same counts for all assemblies, Y-axis is copy number in the assemblies, 2022 T2T, 2020 “Ash”, and 2019 GRC from left to right. X,Y-axes are in log-scale from 1 to 1 million copies; the maximum count found in this genome is ∼20,000. Red dots are mer counts for structural repeats, blue dots are mer counts for coding sequence. Two lines show expected value (r=1, blue line) and observed log-linear regression (r=0.97 for T2T at left, r=0.80 for Ash, r=0.71 for GRC). The T2T graph is very close to the expected 1:1 agreement, showing assembly captures all of DNA content. The earlier GRC assembly has notable drop off at high-copy (right) side of graph, with Ash assembly intermediate, as the regression slopes indicate.

### Corn

**Figure Zm2a,b.**
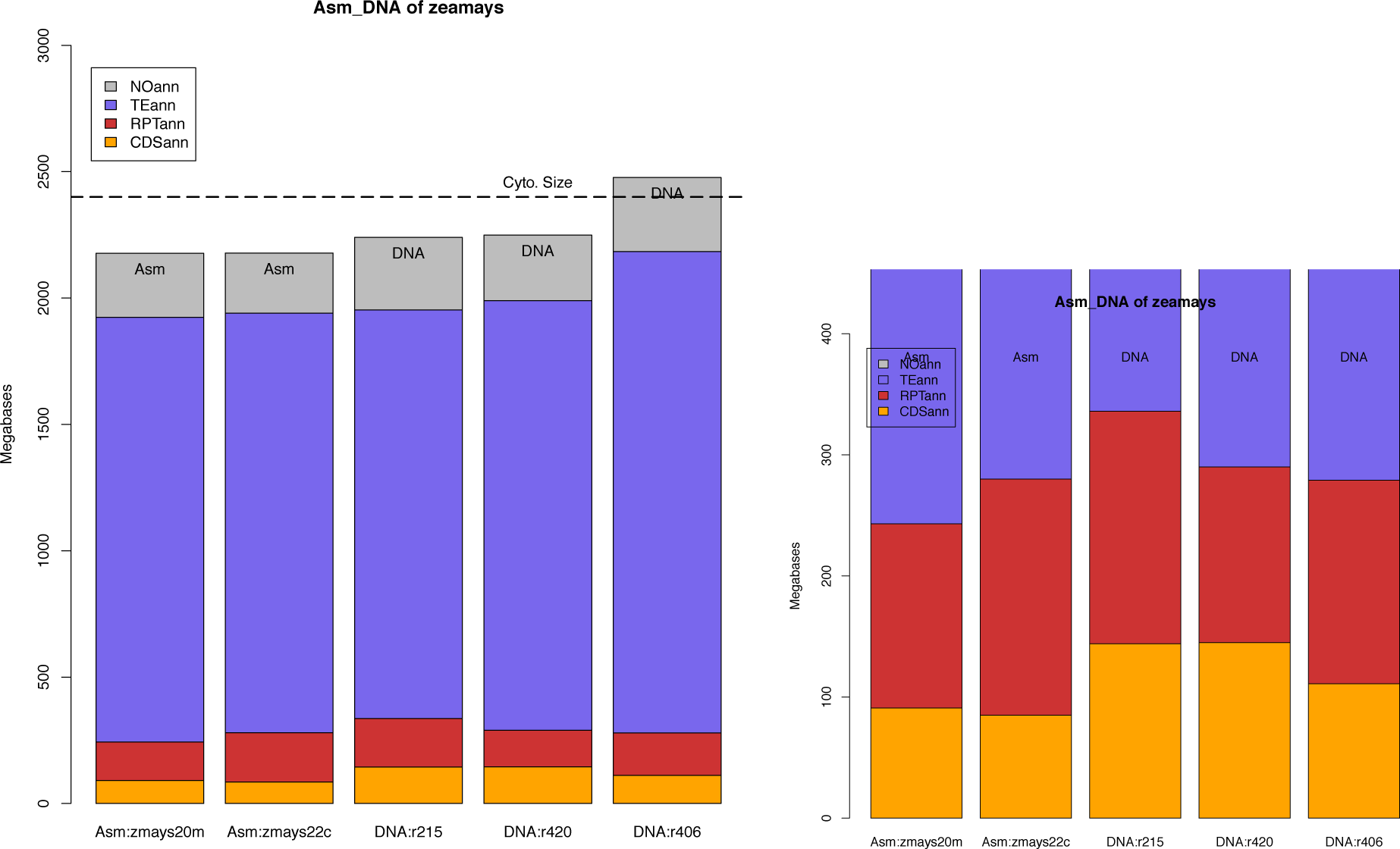
Major contents of *Zea mays* genome assemblies and DNA read sets measured with alignment of DNA on assemblies.

**Figure Zm2c.**
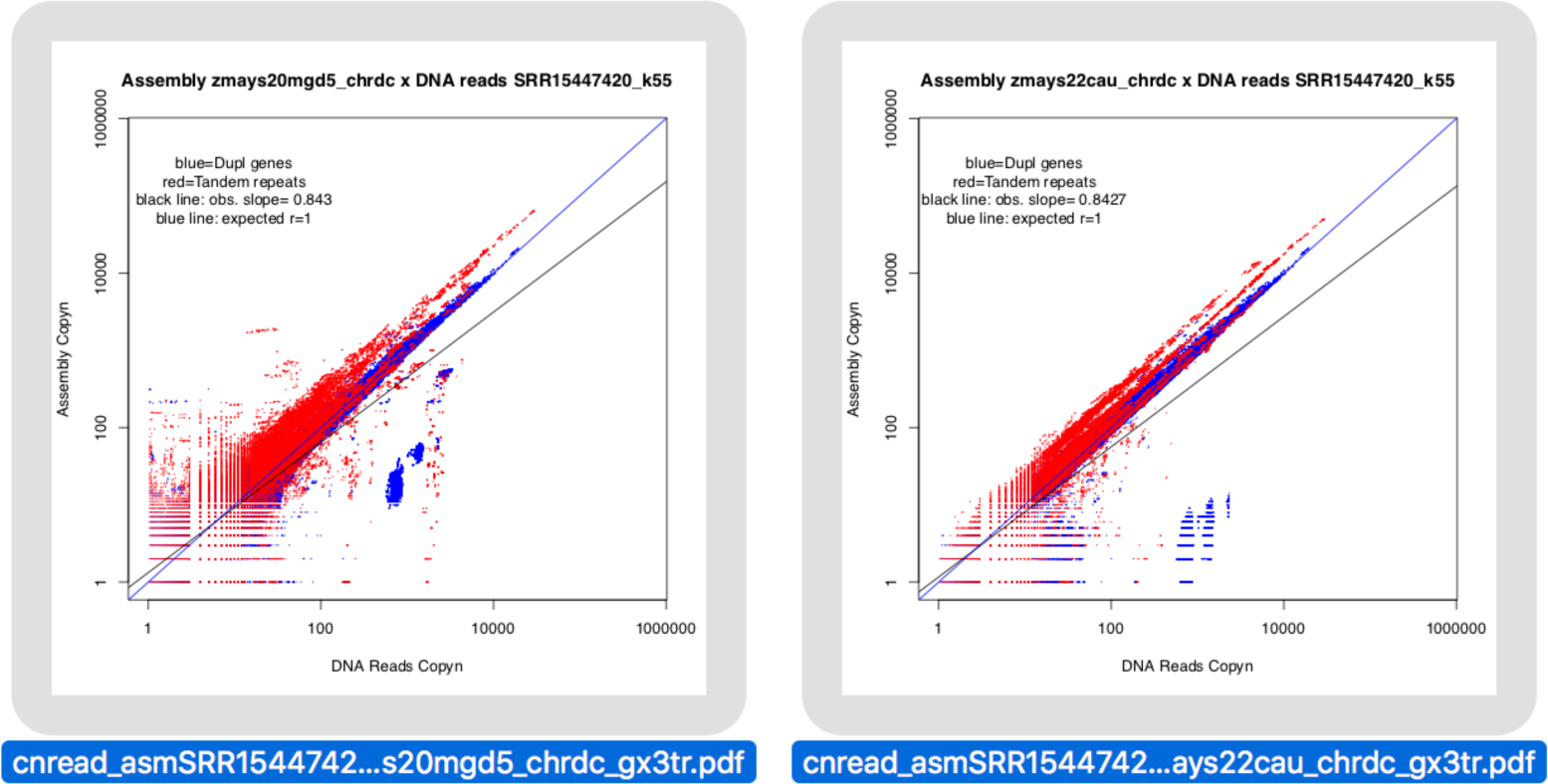
Perfect match counts of 55-mers of high-copy DNA by Assembly for corn, for gene and structural content subsets. Assemblies left to right: Zmays20mgd, Zmays22cau.

### Chicken

**Figure Gg3a,b.**
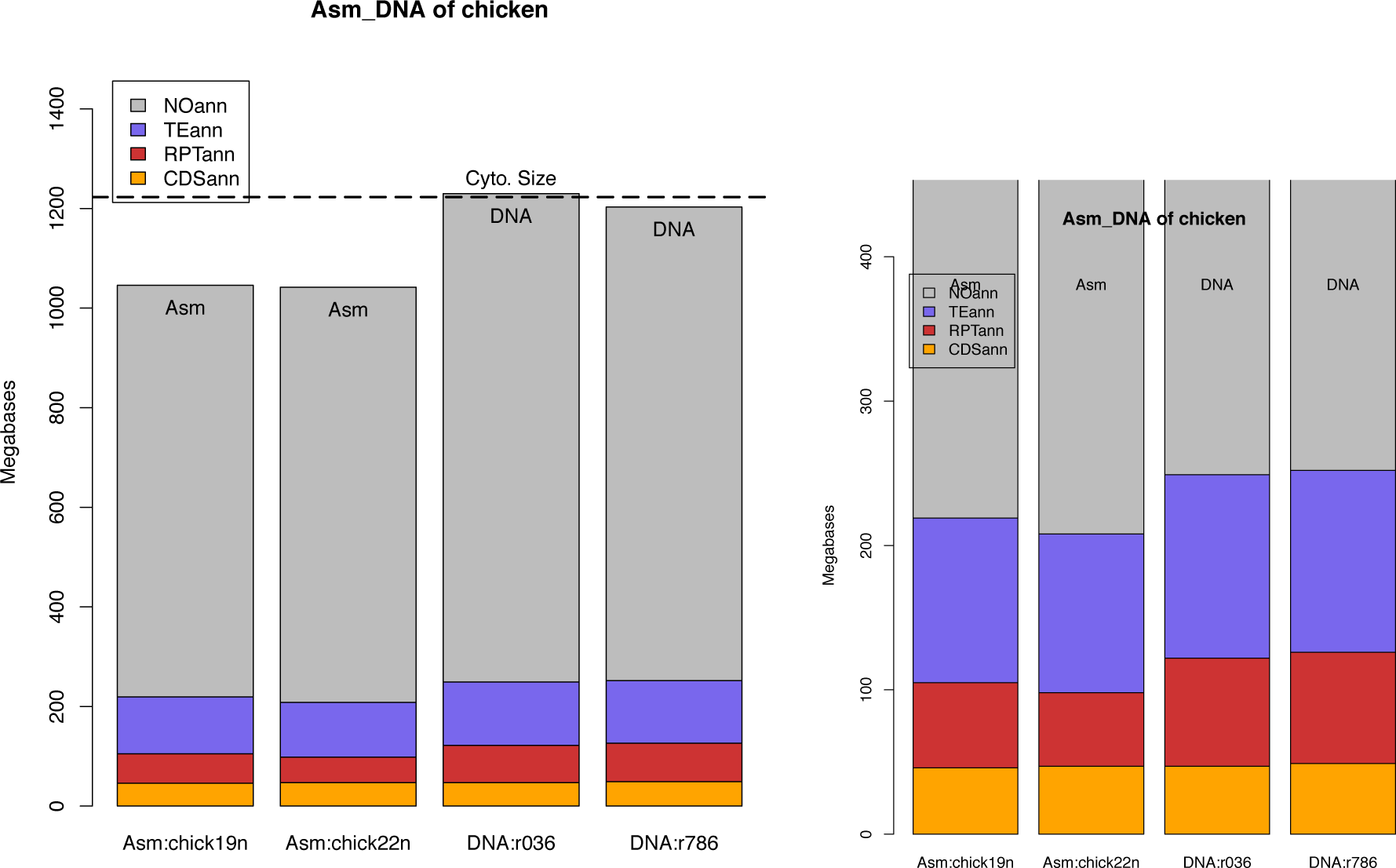
Major contents of *Gallus gallus* genome assemblies and DNA read sets measured with alignment of DNA on assemblies.

**Figure Gg3c.**
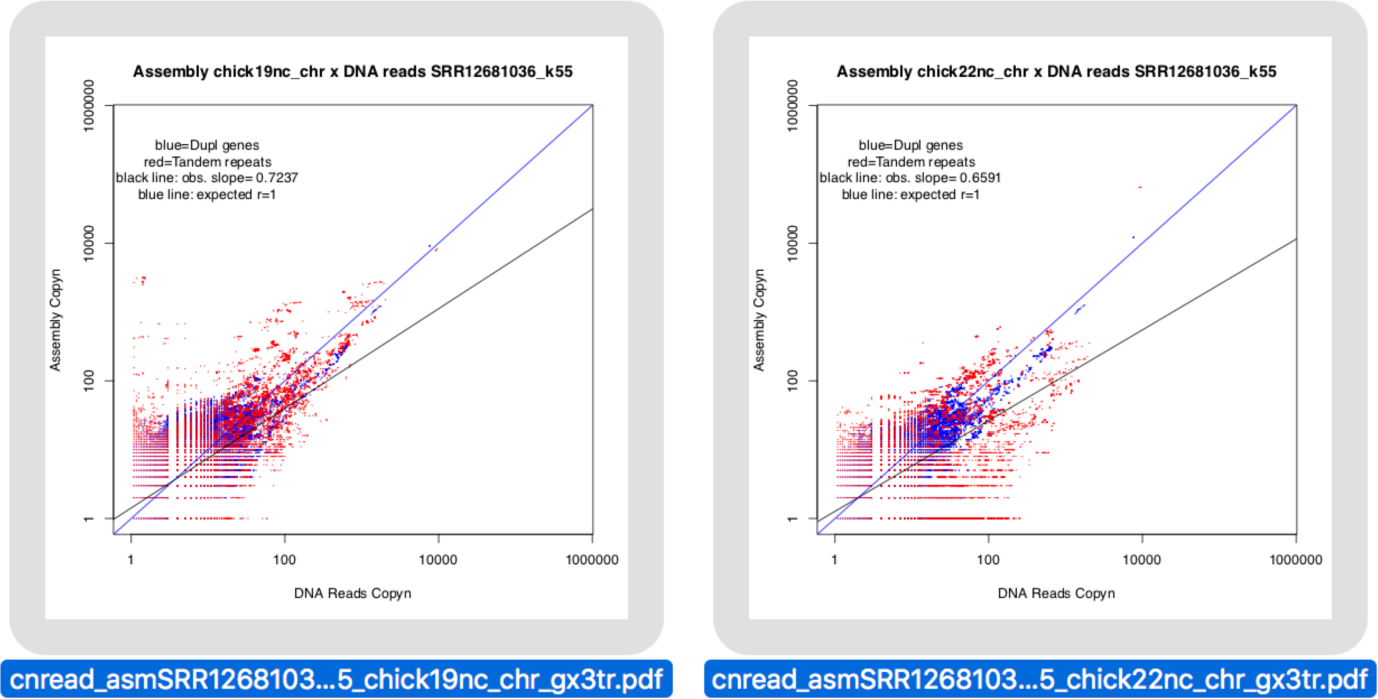
Perfect match counts of 55-mers of high-copy DNA by Assembly, for gene and structural content subsets for *Gallus gallus* genome assemblies. Assemblies from left to right: Galgal19 ncbi, Galgal22 ncbi.

### Ants, flies and lice

The three cases of fireant, blowfly and salmon louse have genome cytometic measures of 600 - 1300 Mb, but down to half that in genome assemblies. A fourth case is for 3 Drosophila with cytometric measures of 250-400 Mb, notably above the 200 Mb of model and other Drosophila.

The blowfly reference genome of 400 Mb, of 2022 compares to 600 Mb cytometric size. DNA size measures with Gnodes yeilds 500-580 Mb, with most of discrepancy in 100 Mb of un-assembled Satellite DNA, 33% of total genome, with 55% Other, 5% Transposons, 6% CDS and 6% other repeats.

**Figure Luc1a,b.**
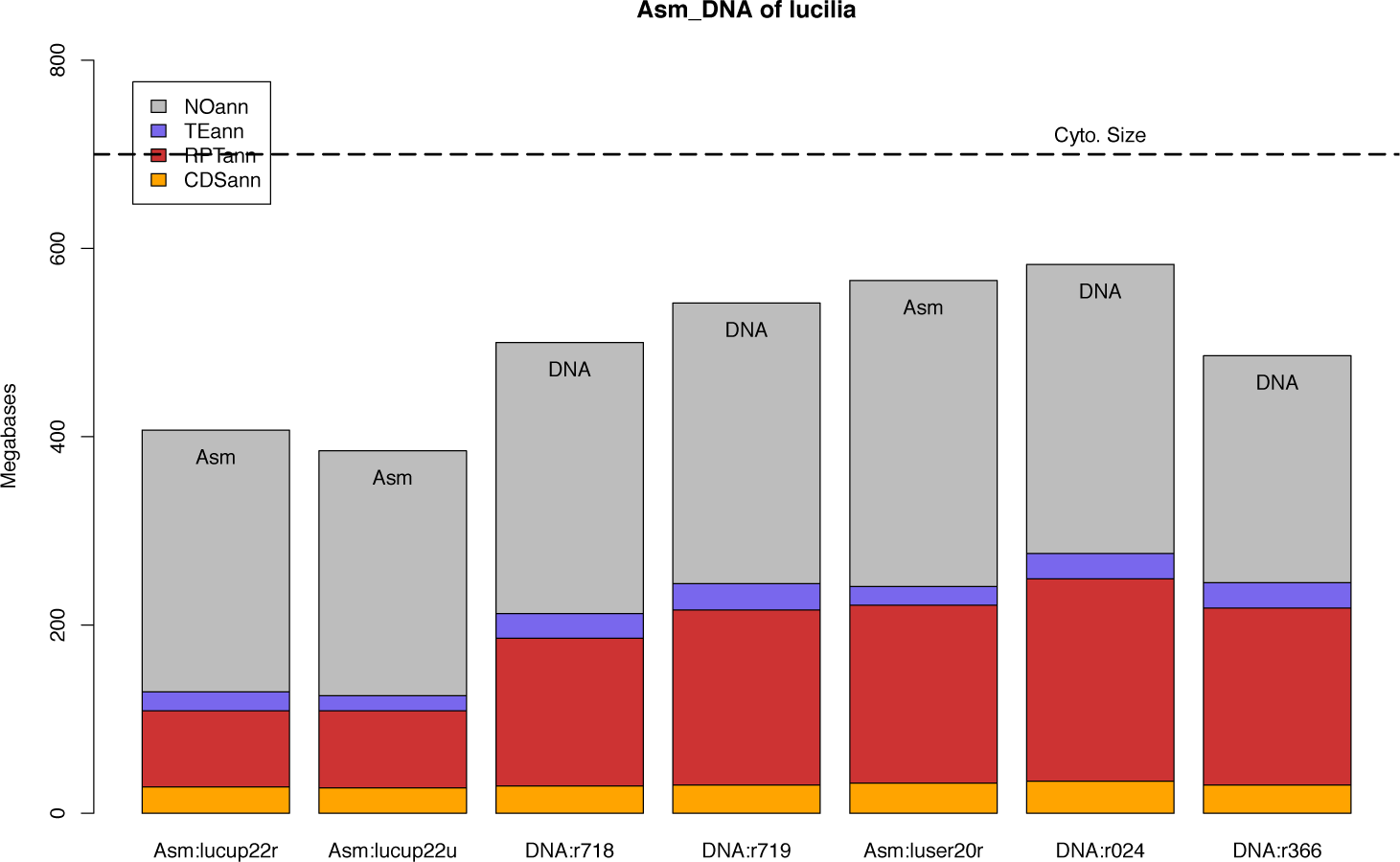
Major contents of blowfly *Lucilia_cuprina* and bottle fly *L. sericata* genome assemblies and DNA read sets measured with alignment of DNA on assemblies. Bar order left-right is L.cup asm1 luccup22ref, asm2 luccup22umel, Dna1 SRR7041718, Dna2 SRR7041719, then L.ser asm1 lucser20ref, L.ser Dna1 SRR10951024, Dna2 SRR13038366.

**Figure Luc1c.**
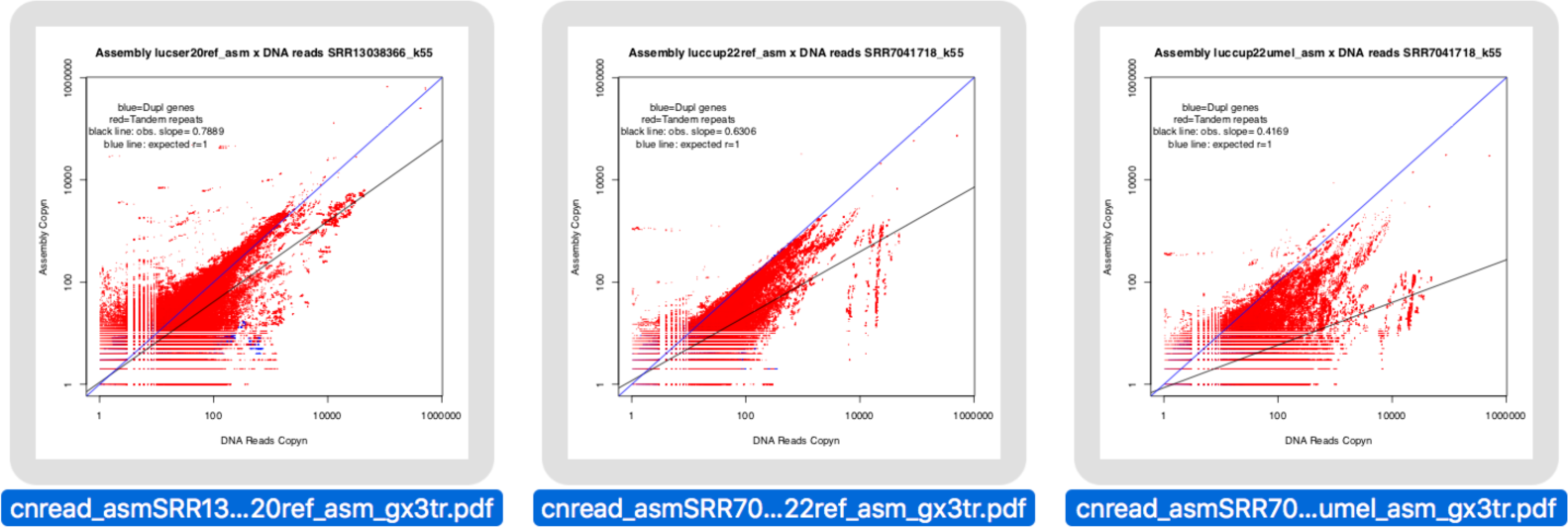
Perfect match counts of 55-mers of high-copy DNA by Assembly, for gene and structural content subsets. Assemblies from left to right, with r slope of asm x DNA match: lucser20ref r=0.79, luccup22ref r=0.63, luccup22umel r=0.42.

Red fire ant DNA similarly contains up to 35% SatDNA of its 600 Mb genomic DNA (measured with cytometry and Gnodes), versus published assemblies of 373 Mb (2021 ref) to 412 Mb (2020 ref). This paper re-assembled the same DNA sample of pacbio with same Canu 2.2 assembler of solin21qmul_asm (384 Mb, Priyam et al. 2021), and obtained 473 Mb in solin23pb2d_asm, including more of the missing SatDNA and some missed gene duplications. The difference here is likely due to use of “purge_haplotigs” on solin21qmul_asm, to remove putative excess heterozygous contigs, but which mistakenly removes valid duplication/repeat contents.

**Figure Sol1a,b.**
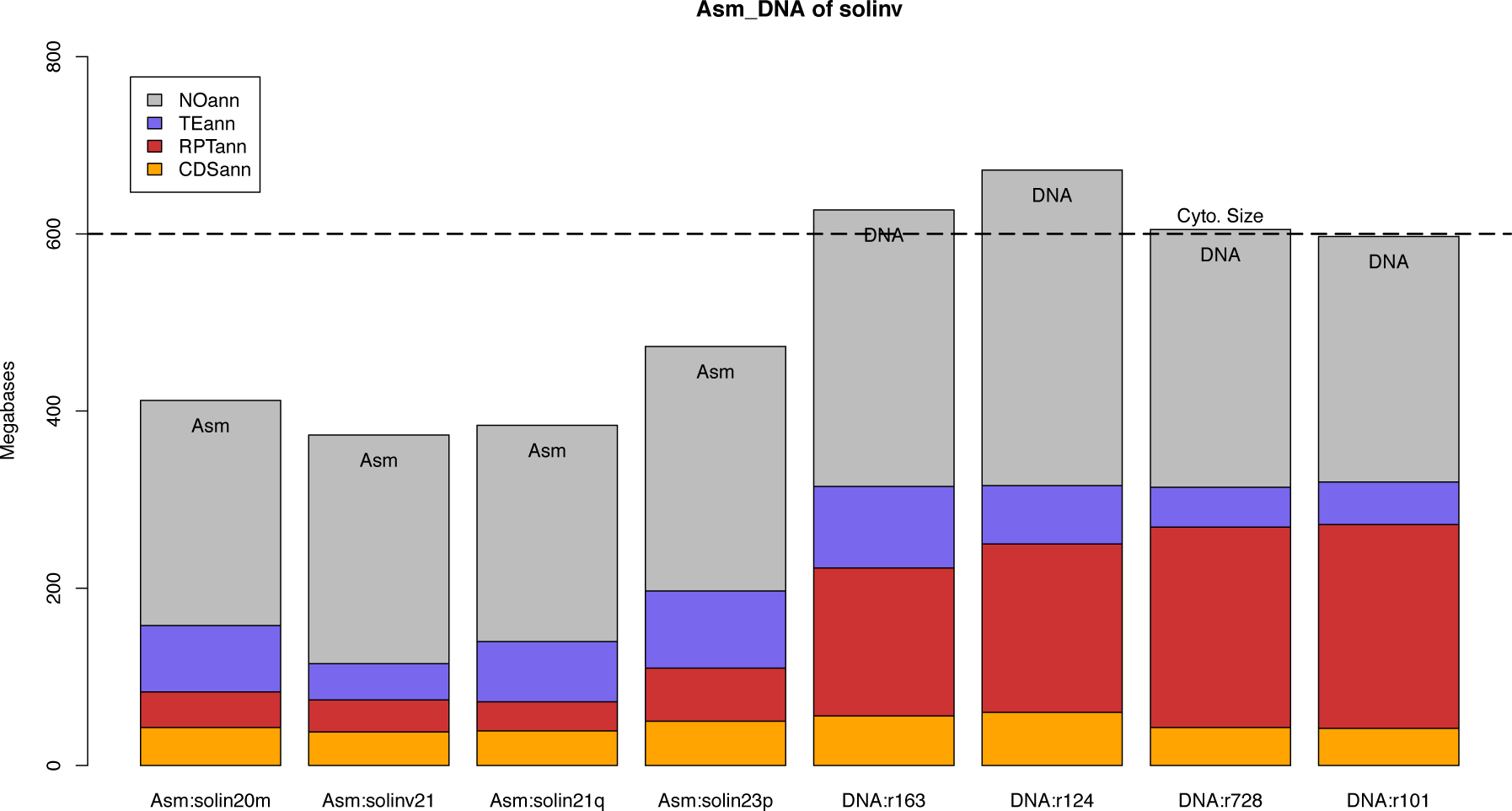
Major contents of fire ant *Solenopsis_invicta* genome assemblies and DNA read sets measured with alignment of DNA on assemblies. Bar order left-right, Asm1: solin20m02sb, Asm2: solinv21ref, Asm3: solin21qmul, Asm4: solin23pb2d, then Dna1 SRR11198163, Dna2 SRR9008124, Dna3 SRR8080728, Dna4 SRR9008101 samples. Dna1 is long-read sample used in Asm3 and Asm4.

**Figure Sol1c.**
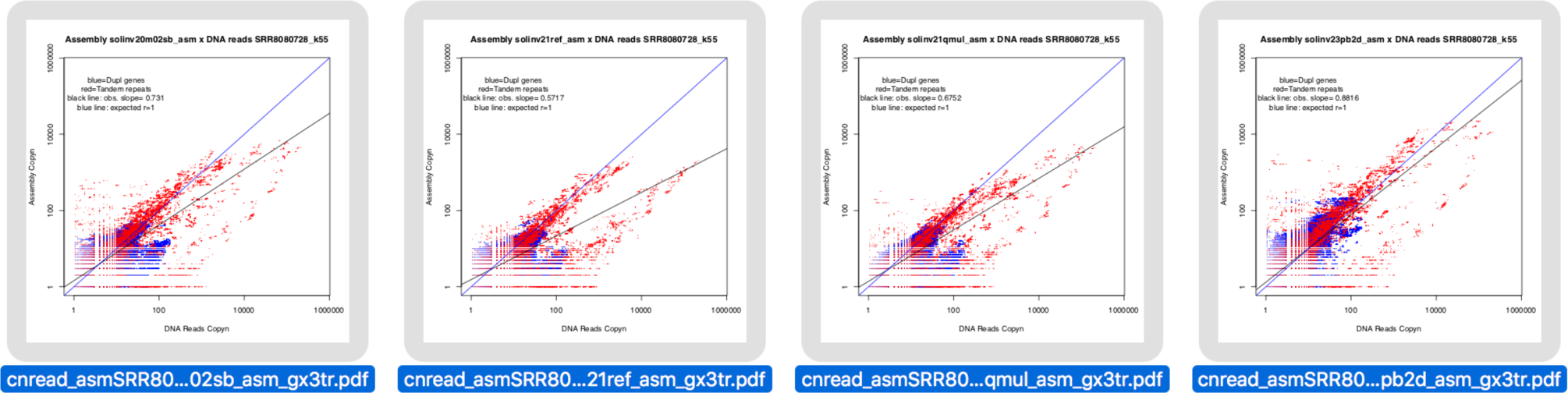
Perfect match counts of 55-mers of high-copy DNA by Assembly, for gene and structural content subsets. Assemblies from left to right, with r slope of asm x DNA match, are Asm1: solin20m02sb r=0.73, Asm2: solinv21ref r=0.57, Asm3: solin21qmul r=0.67, Asm4: solin23pb2d r=0.88 All four assemblies have down-sampled a set of very high copy repeats (10,000 .. 100,000+) to under 100 copies. The r slopes correspond to various levels of down-sampling in more of repeats and gene duplications.

*L. salmonis* crustacean has transposons as its largest major content, 55% of 1000 Mb genomic DNA or 1300 Mb by flow cytometry (Wyngaard et al. 2022), versus 630 Mb of two recent assemblies (lesal21ncbi_chr, lesal21ioa_chr). However, the *L. salmonis* assemblies also under-assemble SatDNA contents by a large order (22 Mb repeats assembled of 80-90 Mb in genomic DNA).

**Figure Les1a,b.**
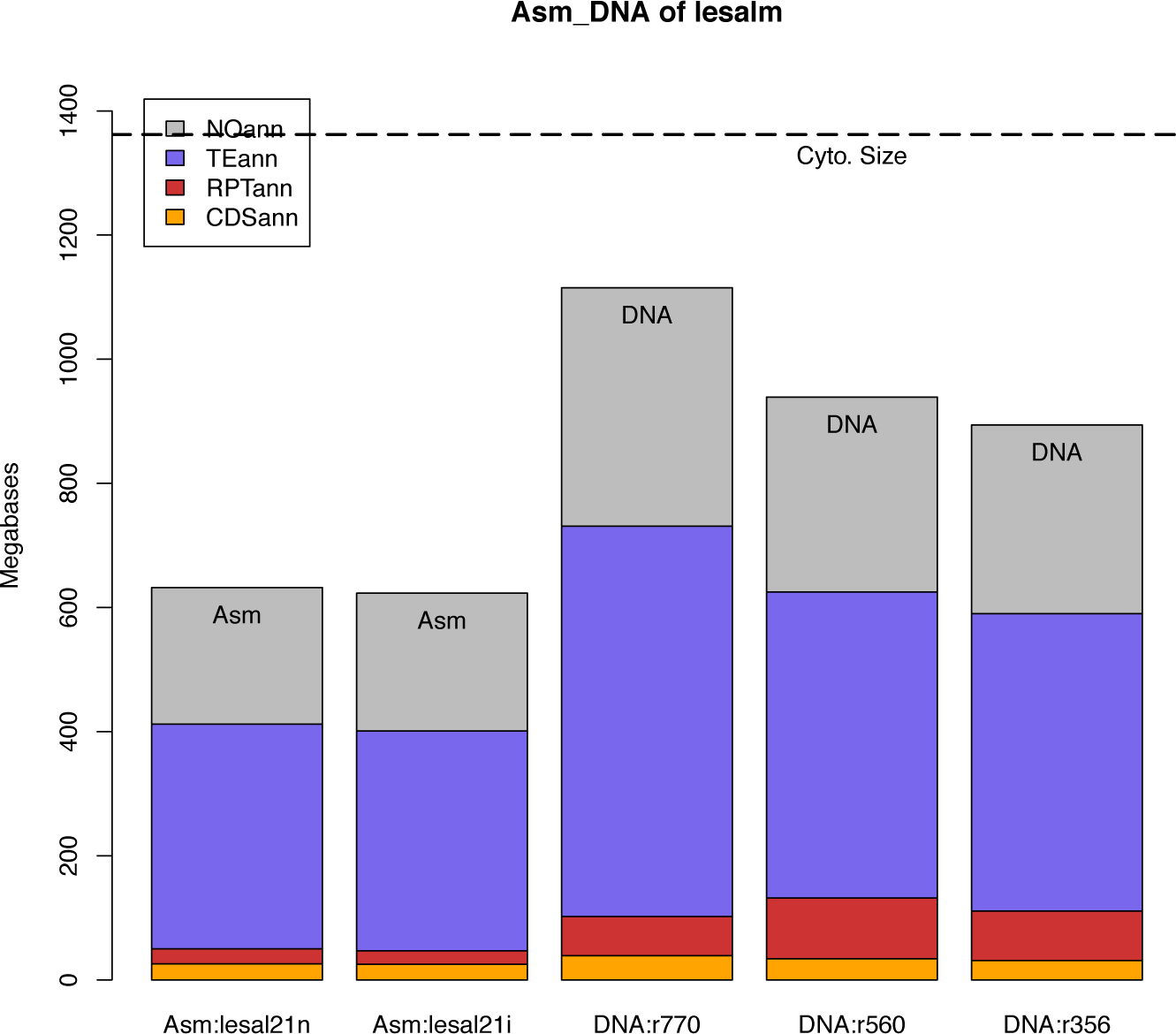
Major contents of salmon louse *Lepeophtheirus_salmonis* genome assemblies and DNA read sets measured with alignment of DNA on assemblies. Bar order left-right, Asm1 lesalm21ncbi, Asm2 lesal21ioa, Dna1 ERR4032770, Dna2 SRR12967560, Dna3 SRR13823356

**Figure Les1c.**
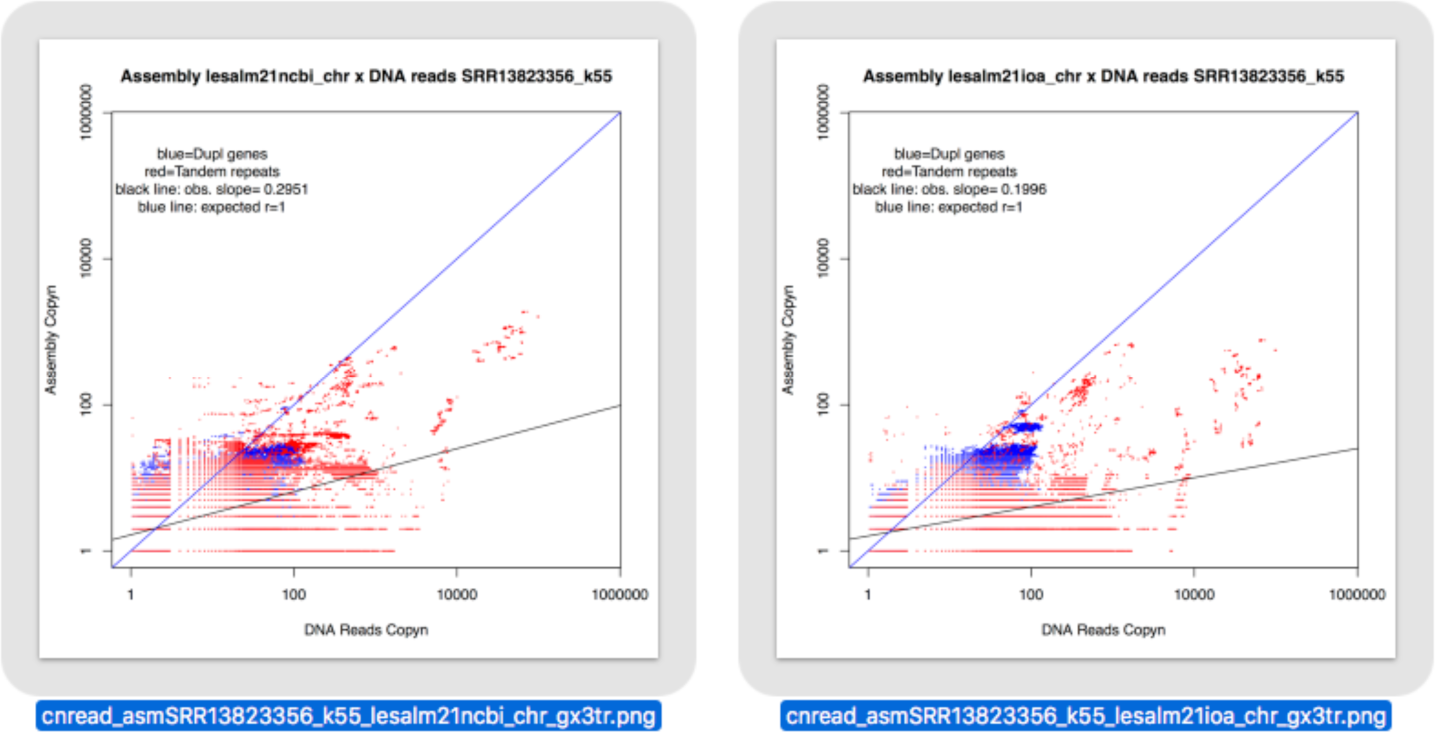
Perfect match counts of 55-mers of high-copy DNA by Assembly, for gene and structural content subsets. Assemblies from left to right, with r slope of asm x DNA match, are Asm1 lesalm21ncbi r=0.30, Asm2 lesal21ioa r=0.20.

For *Drosophila virilis* (cytometry: 350 Mb, Asm: 200 Mb), and *Dro. americana* (cytometry: 300 Mb, Asm: 180 Mb), measurement of DNA samples with Gnodes matches the assembly sizes, i.e. no evidence of large under-assemblies. *D. virilis* current reference assembly, drovir19ref_asm, shows a modest SatDNA under-assembly of 6 to 20 Mb across DNA samples.

For *Drosophila suzukii,* a recently invasive pest fruit fly, the answer is quite different: a 320 Mb size measured with cytometry may be an under-estimate, with DNA sequence measured sizes by Gnodes and K-mer methods above 500 Mb, mostly as repetitive DNA. The early short-read assemblies of 200 Mb are well below measured sizes, with a recent long-read 268 Mb assembly (Paris et al. 2020) still an under-assembly of this genome. Gnodes measures the long-read DNA of this 268 Mb assembly as containing 615-630 Mb, with 100 Mb discrepancies in repeats (SatDNA and other), transposons, and other (un-annotated) portions, plus 30 Mb (2x) of missing gene duplications.

### At-Plant

**Figure At4a.**
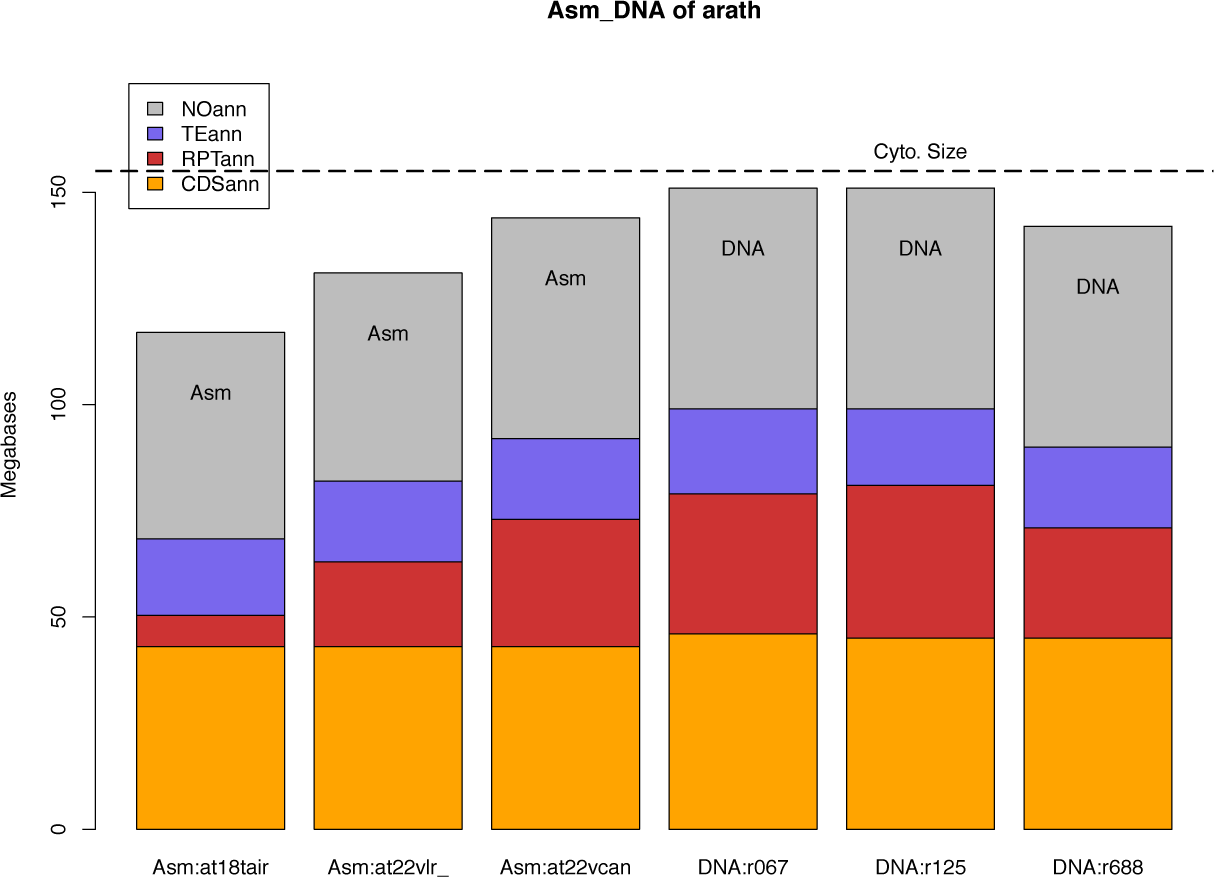
Major contents of *Arabidopsis thaliana* genome assemblies and DNA read sets measured with alignment of DNA on assemblies. This is a stacked bar graph of protein codes (orange, bottom), structural repeats (red-brown), transposons (blue) and other (gray) as portion of the 157 megabase model plant genome. Three assemblies from the left are At18tair (ncbi ref genome of 2018 from TAIR 10), At21vlr (Naish et al 2021), and At22canu (this paper). These differ in size notably in structural repeats, mostly of the telomeric rRNA spans. Three DNA read sets on right, of long and short reads, are left to right, ERR8666067 illumina, ERR8666125 hifi, SRR16841688 hifi, as in Table 3B,C.

**Figure At4c.**
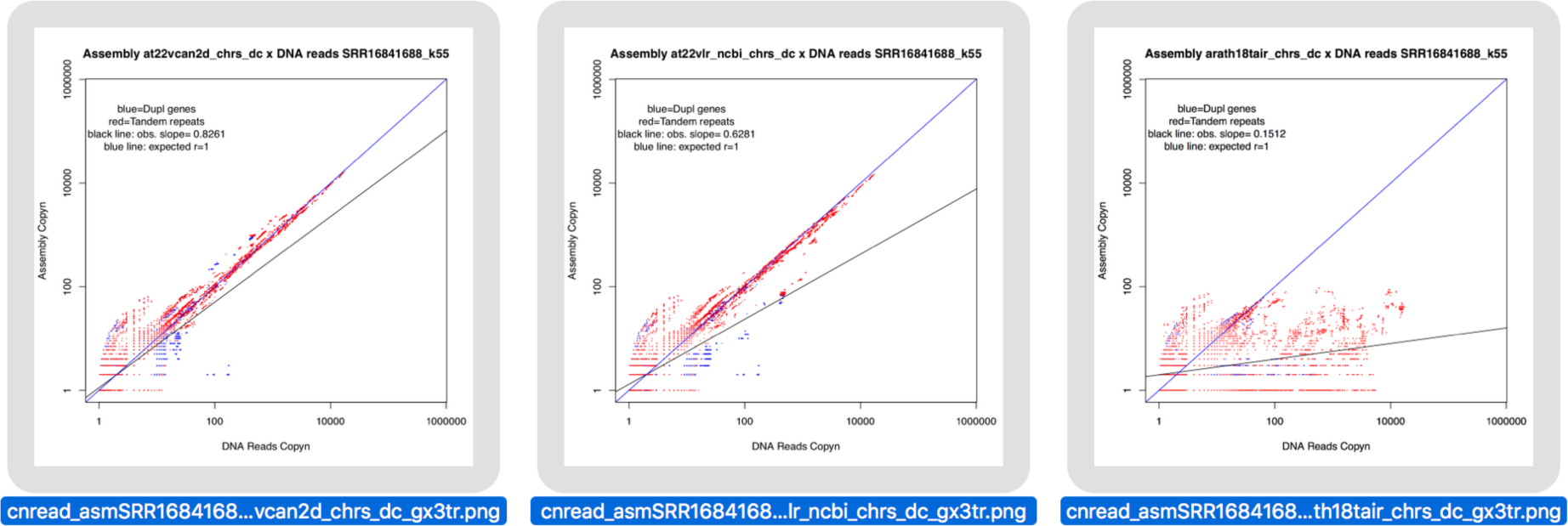
Perfect match counts of 55-mers of high-copy DNA by Assembly for model plant, for gene and structural content subsets. Assemblies from left to right are At22canu, At21vlr, At18tair. X,Y-axes are in log-scale from 1 to 1 million copies; the maximum count found in this genome is ∼50,000. Red dots are mer counts for structural repeats, blue dots are mer counts for coding sequence. Two lines show expected value (r=1, blue line) and observed log-linear regression (r=0.83 for At22canu at left, r=0.63 for At21vlr, r=0.15 for At18tair). The At22canu graph is closest to the expected 1:1 agreement, while still-current reference assembly At18tair is missing much of its high-copy DNA.

Although DNA in Fig At4a measures below an average 157 Mb cytometric size of At species, many more At DNA samples measured with Gnodes do average at this value (see Gnodes #1, #2 docs). The samples here are ecotype Col-0 of the reference assembly, chosen in part due to its smaller genome. Gnodes DNA measures of At ecotypes agree with earlier findings by cytometric measures for longitudinal and other population variation in genome size (Long et al. 2013); these are partly but not entirely related to NOR span changes.

Long read assemblies of At model plant measured here include those of Naish et al 2021, with version 2 of 2022, and 5 other assemblies made by this author from HIFI and ONT DNA reads of that paper (Table At23Asm). Verkko and Flye assemblies are missing significant portions of unique euchromatic genome that all others contain (genes, asm uniq). Canu assembly of ONT-only DNA (300x) suffers from high error rate. at22_Vlr2ncbi of Naish improves some parts but lost some of NOR regions. at22_Canu2HIFI (or At22canu above) and at23_CaNOR6ref are from same Canu2 HIFI assembly, with at23_CaNOR6ref updated for NOR assembly of 2023, and scaffolded with ONT DNA using ntLink (Coombe et al. 2021).

**Table At23Asm.**
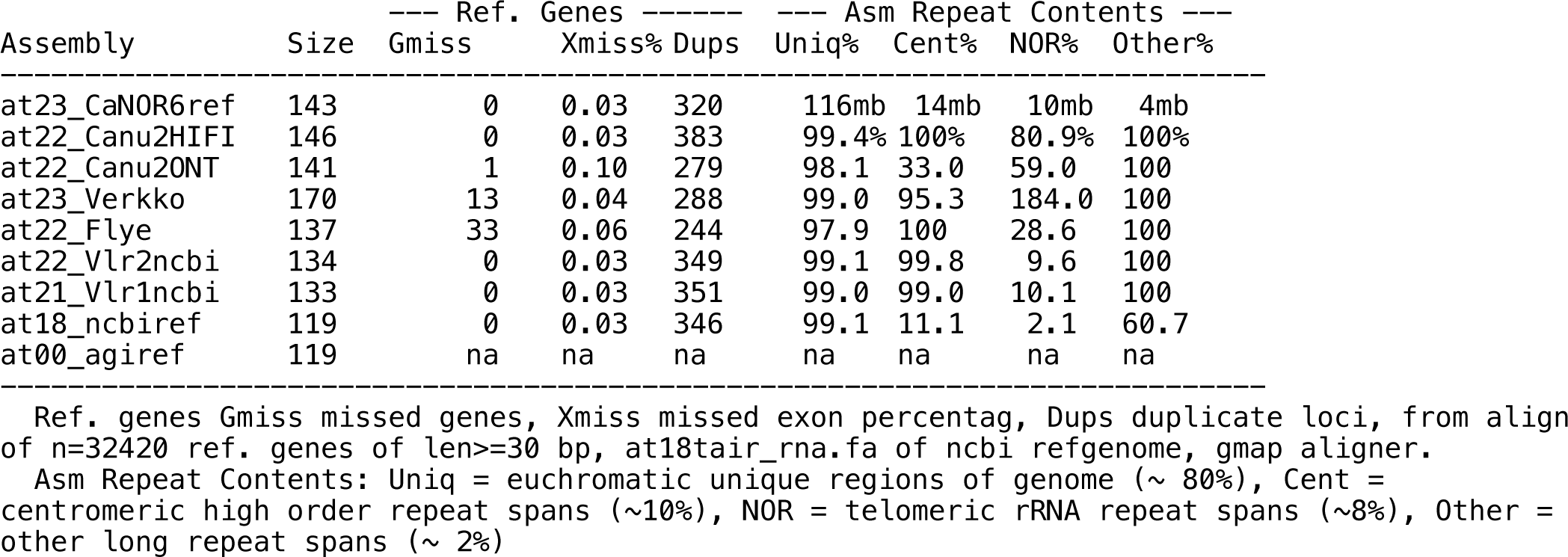
Qualities of At model plant assemblies, relative to complete reference of 2023. These are assembled from Pb HIFI + ONT DNA of Naish et al 2021 (at21_Vlr1ncbi), but for at18_ncbiref (ref TAIR10 of 2018) and at00_agiref (AGI published of 2000).

Verkko, the method developed from work on repetitive human Y-chr (Rhie et al. 2023, Rautiainen et al 2023), came very close to a complete assembly, but for its puzzling failure on some easy unique spans, and NOR spans left as an 184% over-assembly of 1500 contigs. It shares reliance on Canu v2 for Pb HIFI primary data, with the at22_Canu2HIFI and at23_CaNOR6ref assemblies here, and its use of ONT DNA for scaffolding the hifi contigs is similar to that of the ntLink method used for the most complete assembly here, at23_CaNOR6ref. The authors of Verkko did assemble At model plant, using a different PbHIFO + ONT DNA sample set, and obtained somewhat similar near but incomplete chromosomes, missing some of the easy unique spans. A guess is that further optimization or parameter revisions for Verkko would enable it to fully assembly At model plant DNA, and possibly generalize it to work well for other under-assembled species noted here.

### Daphnia waterfleas

**Figure Dap5a,b.**
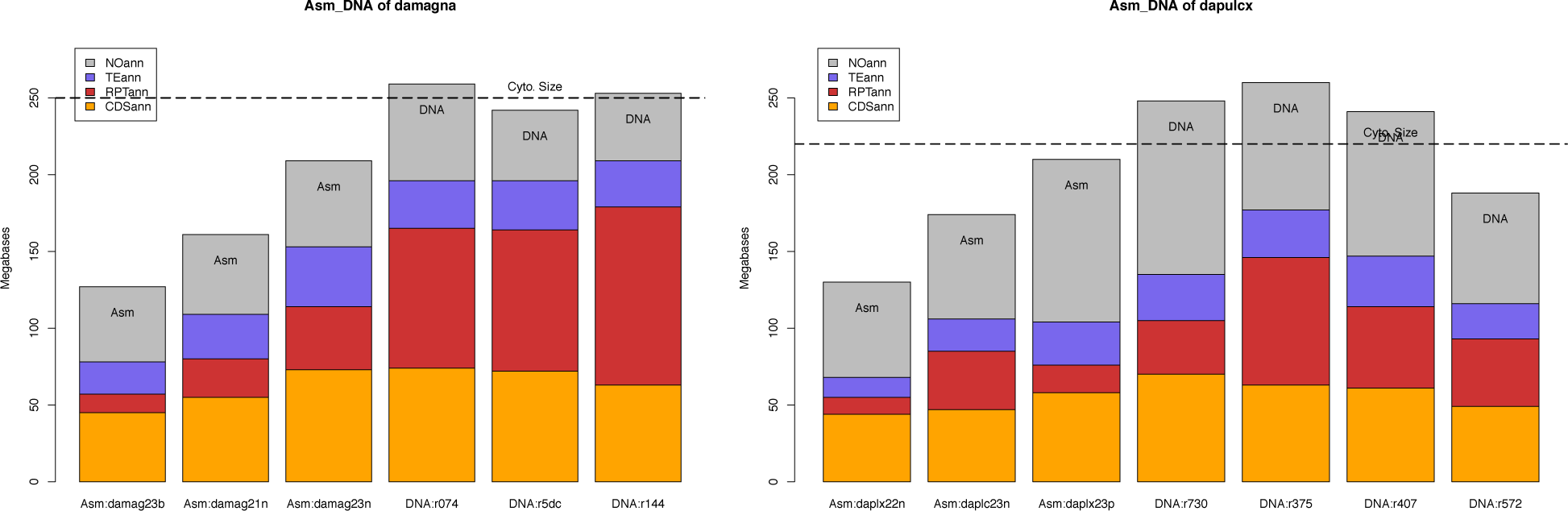
Major contents of *Daphnia magna* (left) and *Daphnia pulex/pulicaria* (right) genome assemblies and DNA read sets measured with alignment of DNA on assemblies. The Y-axis maximum is 250 Mb for both species, with D. pulex average cytometric size of 220 below that of D. magna at 250 Mb. Assemblies of D. magna in Fig 5a, left to right are dmag23bham, dmag21ni7ncbi, dmag23nan4cr7v (this paper), and DNA of SRRnn074, SRRnn5dc, SRRnn144. Assemblies of D. pul. are D. pulex 22 ncbi, D. pulicaria 23 ncbi (r=0.67), and D. pulex23p (this pp), with DNA of SRRnn730, SRRnn375, SRRnn407, SRRnn572. Sibling species D. pulex and pulicaria were measured with DNA of both species, with approx. same results; measurable species content differences are overshadowed by larger artifacts of assembly.

**Figure Dap5c.**
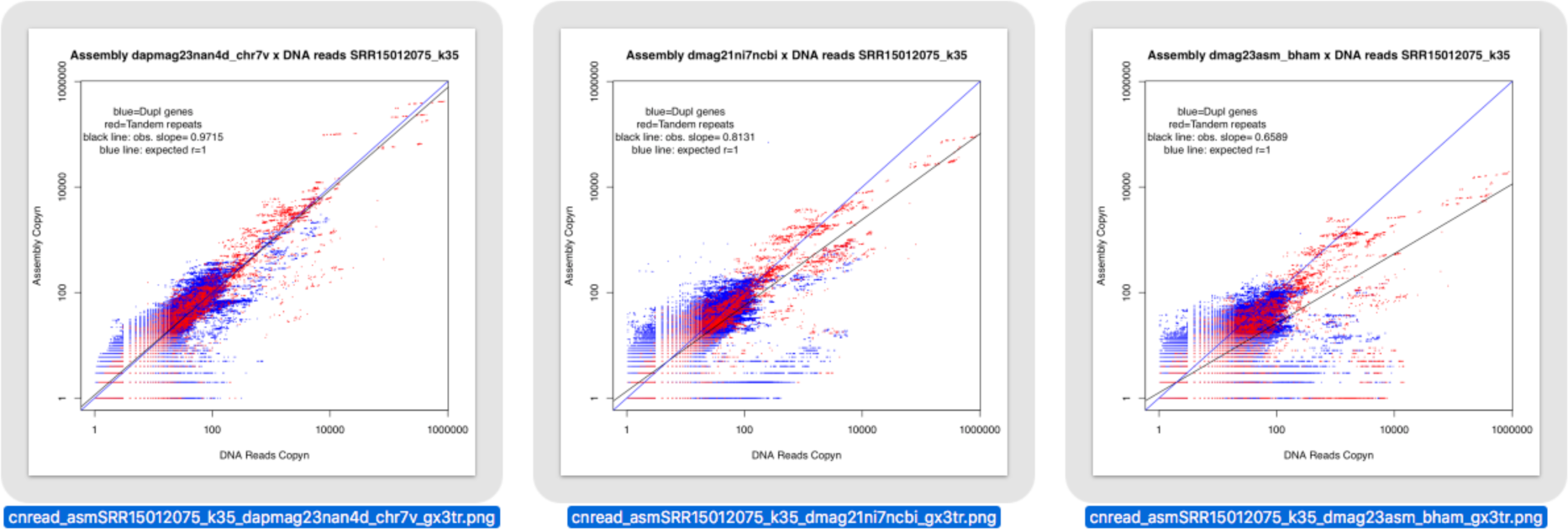
Copy numbers of DNA fragments that are common to DNA reads and assemblies of D. magna. X-axis is mer copy number of DNA-reads, with same values for all assemblies. Y-axis is mer copy number in each assembly, for this log-log plot from 1 to 1 million copies. Two genome content types have mers plotted: duplicated gene CDS (blue dots), and tandem repeats (red dots). Two lines, the equal content line (r=1) and linear regression through observed values (r in table). The three assemblies of D. magna, left to right dmag23nan4cr7v (r=0.97, this paper), dmag21ni7ncbi (r=0.81), dmag23bham (r=0.65), and DNA SRR15012075, described in text. Very high copy number mers, above 10,000 reaching 600,000 copies for the highest repeat copy number of all examined species, are most deficient on assembly Y-axis of recent dmag23bham assembly.

**Figure Dap5d.**
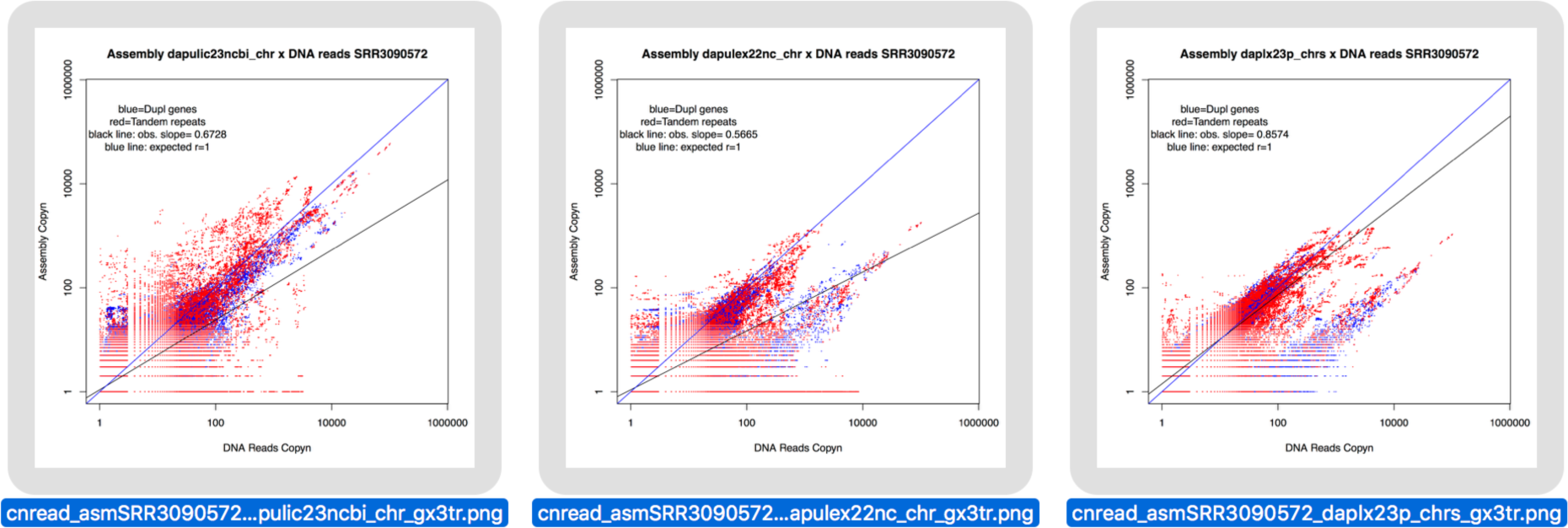
Perfect match counts of 55-mers of high-copy DNA by Assembly for *Daphnia pulex/ pulicaria*, for gene and structural content subsets. Assemblies are D. pulicaria 23 ncbi (r=0.67), D. pulex 22 ncbi (r=0.57) and D. pulex 23p (r=0.86, this paper) by DNA of SRR3090572 (D. pulex).

Copy numbers of DNA fragments (35-mers) that are found in DNA reads are compared to copy numbers of same fragments in chromosome assemblies, for CDS duplicates and satellite repeats, in Figure Dap5c. DNA fragment copy number is the coverage count divided by a constant Cucg, coverage at unique conserved genes. Assembly fragment copy number is the fragment count. The long reads are Nanopore of SRR15012075 used in two of these assemblies, with bacterial DNA removed.

In D.magna, the single highest copy repeat is 70-mer of this form, with duplex and higher order versions spanning 500 Kilobases to Megabases.

>dm21ni074k70c3101895i05147 kc=3101895 == dm21ni074r_circ1.srf.fa AATATACTTGTATTCAATGTGAATACACTGTTATACCATGAGTTTTTACTTCTATATGTAAACTGCATGG

This “dm21ni074r_circ1” 70-mer occurs 3,113,587 times in genomic DNA with 15x coverage depth, or 15 Mb of a 234 Mb genome size. It occurs at lower rates in most assemblies of those DNA sets, 21691 times or 1.5 Mb in the NCBI reference (dmag21ni7ncbi), but 209958 times or 15 Mb in my reassembly of same long-read DNA set SRR15012075, damag23nan4d.

A log-linear regression slope (r) through the DNA read by assembly copy numbers is calculated, with expected slope of 1 for equal copy number, in Fig. Dap5c. Slope below 1 means an assembly has fewer copies than in the genomic DNA. This is inverse (1/r) of xCopy, but meaning and results are about the same (see Table Gndmag below).

| Assembly | $r$ | $1/r$ | xCopy |
| --- | --- | --- | --- |
| dmag23nan4cr7v | 0.972 | 1.03 | 1.16 |
| dmag21ni7ncbi | 0.813 | 1.23 | 1.44 |
| dmag23asm_bham | 0.659 | 1.52 | 1.89 |

Other DNA reads of three other D. magna population samples, Illumina of 120 to 150 bp, produce similar results for genomic DNA by assembly analysis of k-mer fragment copies. This is another way of measuring agreement between contents of DNA reads and assemblies. Gnodes summary above measures by mapping reads onto assemblies, counting coverage or copy number of DNA along assembly spans with annotated contents. There might be a problem for map-measuring very high copy numbers, if mappers are mis-counting this type of DNA. But this perfect-match k-mers method agree with results of imperfect-match alignments including very high copy contents of genomes.

K-mer fragmentation of DNA sequences to short spans, counting each unique mer for reads and assemblies, allows for exact match counts even with very high copy numbers. This is the approach of k-mer genome size estimators, but adds comparison of shared mers between DNA and assemblies, and selection of mers by genome content (CDS, RPT here). There are drawbacks to this method, e.g. inexact matches from artifacts don’t count, and kmer size choice affects results without biological meaning: short k <= 25 lumps together disparate content types, while longer k >= 50 reduces statistical power of this method.

### Summary of Gnodes estimates for *Daphnia magna* genome assemblies x Nanopore DNA

A summary of Gnodes measures for recent Nanopore long-read assemblies of Daphnia magna is given in Table Gndmag. Obs. and Est. are megabases of total (allasm) and components (CDS coding sequences, TE transposons, simple and tandem repeats (RPT and Satellite-type repeats), and NOann, composed of unannotated portions including non-coding genes, introns and intergenic spans that are not TE or RPT. Estimate is that of DNA read contents, mapped to assembly, while Observed is content of assembly. xCopy is the ratio of read DNA span to assembly span (or read coverage depth normalized by Cucg). Cucg= 118 is read coverage depth, measured from unique conserved gene coverage.

Assembly dmag23nan4cr7v has a near 1:1 ratio for assembly x DNA sample contents, only RPT-Satellite content is low (xCopy of 2). This is also at low end of flow cytometry measured genome size of 220-260 Mb. Note that Est. Mb is close to same value across assemblies, as this is measured content of the same DNA sample.

Satellite DNA (or tandem repeats) and gene CDS duplicates (CDSdupx) are two major genome components showing largest difference among assemblies, accounting for ∼90 Mb discrepancy dmag23nan4 and dmag23bham.

**Table Gndmag.**
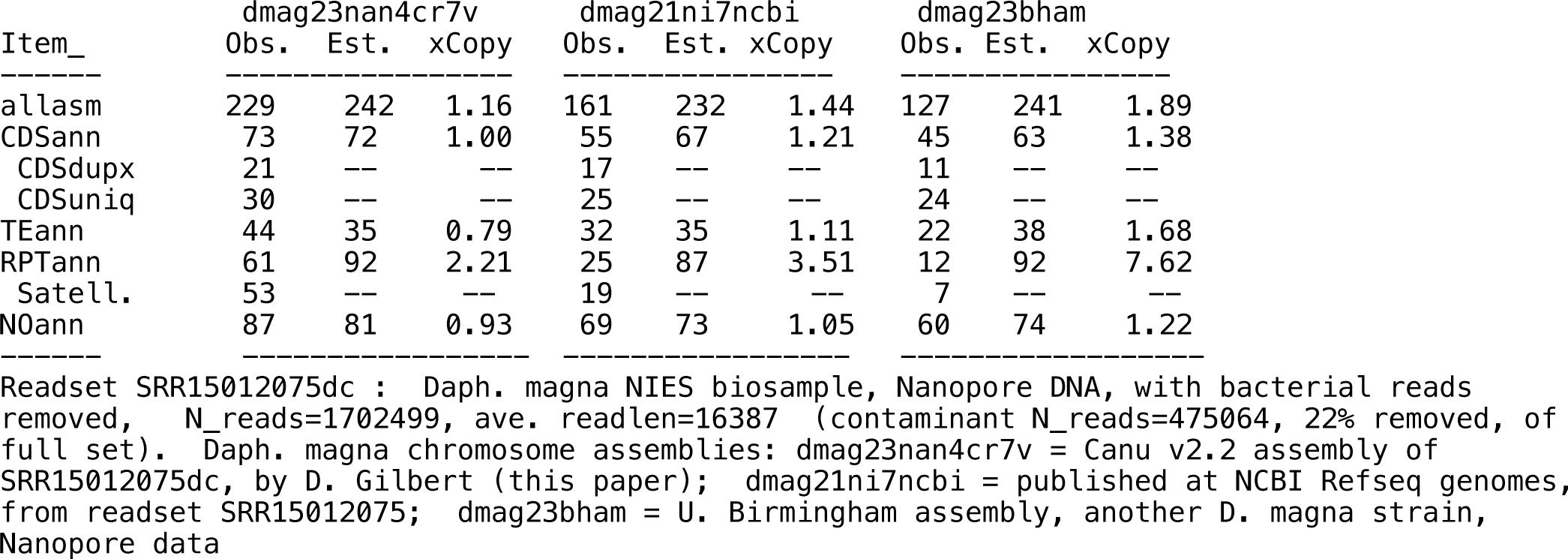
Gnodes summary for 3 Daphnia magna assemblies (gnodes23dm075rmm)

Other DNA reads of three other D. magna population samples, Illumina of 120 to 150 bp, produce similar results, with nearly same estimated contents, and largest xCopy for tandem repeats and duplicated genes. The unique and duplicated gene CDS categories are a subset of the full reference gene set (published 2014), those having unique or duplicated class in 3 separate population DNA samples. Update 2023May20, uses daphmag23arpod_telib.fa, repeat modeler with more TE, tandem repeat trf.tab for SatDNA.

**Table Trd.** Tandem repeats measured in Daphnia and model genome assemblies. Tandem repeat finder (trf) results with TR span counts, % and megabases of TR content.

| Assembly | TR count, TR total bases |  |  |  |  |
| --- | --- | --- | --- | --- | --- |
| damag23nan4d | 9144, 20% | 46.0 Mb | of 238 Mb | asm | *Daphnia magna* |
| damag21ni7ncbi | 4688, 10% | 16.2 Mb | of 160 Mb | asm | " |
| damag23bham | 2618, 4% | 4.7 Mb | of 130 Mb | asm | " |
| dapulic23ncbi | 11884, 12% | 21.9 Mb | of 185 Mb | asm | *Dap. pulicaria* |
| dapulex23pbc | 4575, 3% | 5.5 Mb | of 220 Mb | asm | *Daphnia pulex* |
| dapulex22az | 1890, 2% | 2.5 Mb | of 130 Mb | asm | " |
| dapulex19pa42 | 3102, 2% | 3.0 Mb | of 185 Mb | asm | " |
| daobtusa20 | 7871, 8% | 12.9 Mb | of 168 Mb | asm | *Daphnia obtusa* |
| at-plant | 3059, 10% | 15.1 Mb | of 150 Mb | asm | *Arabidopsis* |
| chick | 13729, 2% | 18.4 Mb | of 1065 Mb | asm | *Gallus gallus* |
| corn | 59838, 2% | 38.1 Mb | of 2182 Mb | asm | *Zea mays* |
| human | 55145, 7% | 214 Mb | of 3117 Mb | asm | *Homo sapiens* |

[revise table .. update w/ DNA measure of HOR, add fireant, others ?]

## Methods of measuring major contents

The major contents of genomes presented in these results are computed with Gnodes, as described in Gnodes#1 (Gilbert 2022). The basic algorithm of Gnodes is (a) align DNA reads to gene and chromosome assembly sequences, recording all multiple and unique map locations, (b) tabulate DNA cover depth at each sequence bin location for multiple and unique mappings, (c) measure statistical moments of coverage depth per item, itemized by categories of gene-cds, transposons and repeat annotations, duplicate and unique mappings, (d) summarize coverage and annotation tables, at chromosome-assembly and gene levels. The statistical population of coverage is non-normal, so that median, average, skew and other statistics are calculated to approach precise and reliable measures.

### Methods data sets

Public SRA IDs of genomic DNA used for each species assembly, as tabled in species23chrdna_gdupstr_sum2b.txt, are provided in suppl. table gnodes23chrasm_dnatable.txt. Genome assembly meta data, including ID, Species, size, GenbankID or Asm URL, Date, are provided in suppl. table gnodes23chrasm_metadata.txt

Genome cytometric and assembly sizes for arthropods (crustacean and insect only), are tabulated from Animal Genome Size Database (AGSD, Gregory, 2023) and NCBI Genomes (2023) reports, in suppl. asm_cytosize_table_arthropods.txt. This table has one row for each NCBI genomes assembly of corresponding species. Columns from include Gsize = cyto measured size, median; EqualGA = equal or DIFF if Asize and Gsize are same or not, by 15%; Class = species class; Mbases, C_value = cyto size measured 1C pg value, and megabases as 978*C_value; Method = FC, FIA, FD or other cytometric measure; MBrange = range and count of cyto measures. Columns from NCBI are Asize and Size (Mb) = assembly size, Assembly Accession = NCBI ID, Scaffolds, Release Date per NCBI table.

Summary table of major contents of assemblies and DNA in animals and plant genomes, calculated with Gnodes are in suppl. species23chrdna_gdupstr_sum2b.txt. Columns are ChrMBob = Chr assembly Megabases, observed, i.e. assembly size; ChrMBes = Genomic DNA Megabases (estimated) from mapping DNA to assembly, measured w/ Gnodes CDSann, CDSbus, CDSdup : size of gene CDS, all annotated, uniq conserved (bus) and duplicated; RPTann, RPTsat : repeat content, and Satellite subset, measured with RepeatMasker, RepeatModeler, TandemRepeatFinder, and SatelliteRepeatFinder; TEann

: transposon content, measured with RepeatMasker, RepeatModeler, or from NCBI reference genome repmask output table; NOann : unannotated as one of above. allasm, dupasm are subset measures of Gnodes, not used here. Last column is name of assembly.DNAsample

### Methods of genome re-assembly

Canu v 2.2 is used for several re-assemblies of published genome DNA and assembly.

Software components of Gnodes are used in reassemblies, of evigene/scripts/genoasm/, include asm_evd_filter.pl : filters extra duplicate contigs of genome assembly using Gnodes DNA depth, gene and other evidence annotations; and asmlink2refchr.pl : uses reference chromosome assembly, and chromonomer software to scaffold contig assembly to chromosome level.

*Arabidopsis_thaliana* re-assemblies include these Canu assemblies using DNA of Naish et al 2021: at22_Canu2HIFI or at22vcan2d_chrs made with Canu v2.2 with PbHifi (SRR16841688), over-assembled duplicate contigs removed using Gnodes asm_evd_filter.pl, then scaffolded to chr-level with asmlink2refchr.pl and at21_Vlr1ncbi assembly.

at22_Canu2ONT made with Canu v2.2 with Nanopore (SRR16832054), over-assembled duplicate contigs removed using Gnodes asm_evd_filter.pl, then scaffolded to chr-level with asmlink2refchr.pl and at21_Vlr1ncbi assembly.

at23_CaNOR6ref, is built using contigs of at22vcan2d, updated with published NOR assemblies of Fultz et al. 2023, and scaffolded with ntLink and ONT DNA to chromosome level.

at23_Verkko, is made with Verokko with default options, and the PbHifi and Oxford Nanopore DNA indicated above. at22_Flye is made with flye assembler.

*Daphnia_magna* reassembly damag23nan4dc, size=238, date=2023, in supplement gnodes_newasm/ daphnia_magna23chrs/, is built with Canu v2.2 with Nanopore DNA (SRR15012075), with bacterial reads removed, over-assembled duplicate contigs removed using Gnodes asm_evd_filter.pl, then scaffolded to chr-level with asmlink2refchr.pl and reference scaffolds of dmag21ni7ncbi.

*Daphnia_pulex* reassembly dapulex23pb_chr, size=222, date=2023, in supplement gnodes_newasm/ daphnia_pulex23chrs/, is built with Canu v2.2 with Pb-lofi DNA (SRR10160730).

*Drosophila_suzukii* reassembly drosuzi23pb3e, size=278, date=2023, in supplement gnodes_newasm/ dro_suzukii23asm/, is built with Canu v2.2 with Pb-lofi DNA (SRR10716nnn in 48 parts).

*Solenopsis_invicta* reassembly solin23pb2d, size of 478, dated 2023, in supplement gnodes_newasm/ fire_ant23asm/, is built with Canu v2.2 with Pb-lofi DNA (SRR13610044).

### Methods of major contents results

Methods of major contents results, Figures M1,2,3 and Species detail figures are drawn from Gnodes summary tables of megabases of content, for observed assembly and DNA estimated from unit coverage depths. See suppl. species23chrdna_gdupstr_sum2b.txt

Methods of Figure I1, Genome sizes for animal and plant species, as Assembly / Cytometric size percentage in relation to cytometric sizes. NCBI Genomes (NCBI 2023) provides assembly sizes, median value for multiple assemblies, from year 2020 to 2023, many of long-read DNA methods. Animal Genome Size Database (Gregory 2023) and Plant DNA C-values Database (Leitch et al 2019) provide the most recent cytometric sizes of the same species These data used in figure are tabulated in suppl. table eukaryote_genoacsizes1a.txt.

Methods of Tables 1 and 2, the results for 12 species: Genome sizes from cytometric, DNA samples and assemblies are drawn from cited animal and plant cytometric databases, NCBI genomes database, and Gnodes measures of the DNA sample contents.

Methods of Figure Hs1c and perfect match counts of high-copy DNA by Assembly for other species. Assembly contents of gene coding sequence and repeats were extracted from assemblies using the Gnodes annotations of these contents. Bi-variate k-mer histograms are calculated with KMC, by reducing both the assembly contents and DNA samples to k-mers of chosen size, then finding the intersection of k-mer frequencies for both variates. The plots show the frequency of mers in these bi-variate histograms, at the intersections of DNA and assembly contents. The X-axis is copy number in one DNA read set, with the same counts for all assemblies, Y-axis is copy number in the assemblies, with X,Y plotted in log-scale.

Methods of Figure T1. Animal and plant genome sizes, assembly relative to DNA contents, with missing major contents. Genomes are ordered big to small as in Table 1 on x-axis, with addition of *Beta_vulgaris* (sugar beet 743 mb), *Theobroma_cacao* (chocolate tree 431 mb), *Drosophila_americana* (190 Mb) that lack full repeat content analysis. Boxplot bars (bisque color) represent sizes from NCBI assemblies published at/after 2020, mostly long-read DNA, as percentage of the median measured DNA sizes on y-axis (60% to 100%). Missing contents of CDS, RPT, TE and Other are represented as Assembly/DNA percentage differences, stacked bars to right of genome boxplot, data as in figures M1,2,3. Data of suppl. table allspp_ncbiasmsize2020tab.txt

Methods of Table 3A. Re-map rates and contents of long-read unplaced spans; from Gnodes#2 table S7b_longread_unplacedbases. Unplaced sequence spans, 150 bp or longer, from long reads that partly map to chromosomes were extracted and re-mapped to assemblies, using both blastn and minimap2. Bases that re-align to assembly are tabulated, including their annotated assembly content types.

Methods of Table 3B. Removal of true duplications by popular heterozygosity dedup software, purge_dups and purge_haplotigs, for human and At plant on chromosomes with low and high Satellite DNA content. Input data for both purge_dups and purge_haplotigs are assemblies of At_plant full chromosomes (at23vc2nl6b_nor_chrs) and human T2T chromosomes that are split to 50kb parts. Input source DNA for programs are long read Pacbio Hifi DNA of At_plant SRR16841688 (Col-0 assembly ecotype), and human HG002 male son of SRR18244889+890. The software programs are run with default options on these inputs. Results are the tabulation of parts removed by these software, for Chr1 and Chr2 of At_plant, and human ChrX,Y (haploid), Chr8,9 (diploid in DNA), to show effects on chromosomes with low and high SatDNA content.

Methods of Table 3C. Genome size estimates from histograms of k-merized DNA samples, full (Hk-full) and default truncation at frequency 10,000 (Hk-deflt), for At plant, Daphnia magna, fire ant, salmon louse and corn plant. Sizes are estimated from histograms by GenomeScope, findGSE, covest-repeats. Gnodes estimate from DNA sample is also shown. DNA samples are reduced with k-mer tabulator KMC (ref) to produce a frequency histogram used by k-mer genome size estimators of GenomeScope (v2), findGSE, and covest. K-mer settings were varied to test effects, from 29 to 55. The histogram calculation of KMC and Jellyfish have high-copy cut-off options, which are varied for this test: full uses a high-cutoff of 999999999, and the default software cutoff (10,000 for both?). k-mer histograms produced using Jellyfish results in same histogram as KMC given same k-mer and cutoff settings. For the model plant, non-nuclear DNA amounts measured with Gnodes are removed from k-mer GSE values. For Daphnia magna Nanopore SRR15012075 DNA sample that contains bacterial DNA, the sample was de-contaminated removing reads without alignment to the assembled D. magna chromosomes. No significant contamination was noted in the other species DNA samples. Arabidopsis thaliana DNA samples are ERR8666067 illumina, ERR8666125 hifi, SRR16841688 hifi, with kmer values 29 and 55. Chloroplast is removed from Hk-full with Gnodes measure: At1 28 mb, At2 24 mb, At3 14mb. Daphnia magna DNA SRR15012075 nanopore-decon (minus bacteria), with k=35. Sol. invicta DNA is SRR8080728, with k=55. Lep. salmonis DNA is SRR13823356, with k=55. Zea mays DNA is SRR15447420 with k=55.

## Acknowledgements

The following provide shared computational resources in support of this work: Indiana University Research Computing, Extreme Science and Engineering Discovery Environment (XSEDE) then Advanced Cyberinfrastructure Coordination Ecosystem: Services & Support (ACCESS), supported by National Science Foundation, including Jetstream (jetstream-cloud.org) and San Diego Supercomputer Center.

## Notes

http://eugenes.org/EvidentialGene/other/gnodes/gnodesdoc/

### Competing Interest Statement

The authors have declared no competing interest.

